# Translational Control of TFEB and Autophagy via eIF5A Rejuvenates B Cell Immunity

**DOI:** 10.1101/360503

**Authors:** Hanlin Zhang, Ghada Alsaleh, Jack Feltham, Yizhe Sun, Thomas Riffelmacher, Philip Charles, Lisa Frau, Zhanru Yu, Shabaz Mohammed, Stefan Balabanov, Jane Mellor, Anna Katharina Simon

## Abstract

Failure to make adaptive immune responses is a hallmark of aging. In particular reduced B cell function leads to poor vaccination efficacy and a high prevalence of infections in the elderly. However, the molecular mechanism underlying immune senescence is largely unknown. Here we show that autophagy levels are specifically reduced in mature lymphoid cells, leading to compromised memory B cell responses in old individuals. Spermidine, a naturally occurring polyamine metabolite, induces autophagy *in vivo* and rejuvenates memory B cell responses in an autophagy-dependent manner. Mechanistically, spermidine post-translationally modifies the translation factor eIF5A, which assists the synthesis of TFEB, a key transcription factor of autophagy. Spermidine is depleted in the elderly, leading to reduced TFEB expression and autophagy. Replenishing spermidine restored this pathway and improved the responses of old human B cells. Taken together, our results reveal an unexpected autophagy regulatory mechanism mediated by eIF5A at the translational level, and this pathway can be harnessed to rejuvenate immune senescence in humans.

## INTRODUCTION

Immune senescence is characterized by the failure of lymphocytes to respond adequately to infection, malignancy, and vaccination. Infectious diseases are still the fourth most common cause of death among the elderly in the developed world. During a regular influenza season, about 90% of the excess deaths occur in people aged over 65 (Yoshikawa, 2000). Immune responses to vaccines are known to be particular ineffective in the elderly population (> 65 yrs), and yet some vaccines, such as influenza, are primarily given to the elderly. A major correlate of protection for vaccinations is the specific antibody titer generated by long-lived plasma B cells. With a life span of up to several decades, long-lived lymphocytes are particularly prone to accumulate waste over time. Autophagy recycles unwanted cytoplasmic material. Lymphocytes deficient in autophagy are unable to generate adequate responses, in particular long-lived lymphocytes, *i.e.* memory T and B cells or plasma B cells (Chen et al., 2014; Pengo et al., 2013; Puleston et al., 2014; Xu et al., 2014). Reversing or halting immune aging would open opportunities to improve management of age-related morbidities and have a major impact on the health status of society. However, little is known about how immune senescence can be reversed.

Genetic activation of autophagy extends lifespan in mice (Fernandez et al., 2018; Pyo et al., 2013). However, only a handful of autophagy-inducing drugs have been shown to reverse aging, one being rapamycin, an inhibitor of mTOR (Bjedov et al., 2010). Due to the unwanted effects of mTOR inhibition, there is a need to better understand mTOR-independent control of autophagy for drug development. Furthermore, to test anti-aging drugs, biomarkers that can be measured in blood are critical. As an endogenous polyamine metabolite that declines with age, spermidine may be key in controlling cellular aging via autophagy (Eisenberg et al., 2009). Previously we found that spermidine reverses aging of the memory T cell response (Puleston et al., 2014) and Eisenberg *et al.* rejuvenated cardiac function with spermidine (Eisenberg et al., 2016), both in mice, in an autophagy-dependent manner. Here we investigated whether spermidine is able to rejuvenate long-term B cell responses, the main correlate of protection for vaccination. We found an autophagy-dependent improvement of old B cell responses by spermidine, indicating that it has rejuvenating effects across different immune cell types. We further identified the target of spermidine and the mechanism by which it regulates autophagy. Using ribosome profiling, proteomics and biochemical assays, we show that the post-translational modification (hypusination) of eIF5A by spermidine regulates both protein synthesis and autophagy in primary B cells. Due to the rigid structure of proline, triprolines slow down translation and require hypusinated eIF5A to form peptide bonds more effectively (Dever et al., 2014). We find this is specifically required for the synthesis of the autophagosomal/lysosomal master regulator TFEB (Settembre et al., 2011), which is a short-lived protein containing one triproline motif in mouse and two in human. Moreover this pathway is considerably down regulated in peripheral blood mononuclear cells (PBMCs) from the elderly, making its components suitable biomarkers. Importantly spermidine improves hypusination of eIF5A, TFEB protein expression, autophagic flux and antibody titers in old human B cells, which has direct translational relevance.

## RESULTS

### Spermidine Induces Autophagy in vivo but does not Affect Hematopoiesis in Old Mice

Previousy, we demonstrated that autophagic flux is decreased with age in human and murine T cells (Phadwal et al., 2012; Puleston et al., 2014). Here, we first investigated the extent of the decrease across bone marrow progenitors and splenic mature hematopoietic cells, with a method of quantifying endogenous LC3-II, a marker for autophagosomes, by flow cytometry that we adapted for primary hematopoietic cells (Cossarizza and al, 2017). To measure active autophagic flux, we used bafilomycin A1 (BafA1) which is a lysosomal V-ATPase inhibitor that prevents acidification of lysosomes and their degradative functions, thereby causing accumulation of autophagic substrates including LC3-II. While autophagic flux was mildly reduced in hematopoietic stem cells (HSCs), it was significantly decreased in B and T cells from old mice as compared to young mice (Figure 1A, gating strategy in S1A/B). Treatment of mice for 6 weeks with spermidine increased autophagic flux in most hematopoietic cell types tested (Figure 1A). We then tested if the *in vivo* administration of spermidine alleviates typical hallmarks of hematopoietic aging, including expansion of phenotypic HSCs, myeloproliferation, and lymphopenia (Henry et al., 2011). We observed a significant increase of phenotypic HSCs and Lin^-^Sca1^+^cKit^+^ cells (LSKs) in old mice, which spermidine administration did not affect (Figure 1B and S1C). A myeloid-biased phenotype, including increased myeloid cells and more significantly diminished mature B and T cells, was found in spleen of old mice (Figure 1C and S1D). Consistently, multiple bone marrow myeloid progenitors are expanded (Figure 1D and S1E/F). Bone marrow pro-B cells (CD43^+^) also mildly accumulate (Figure 1E/F and S1G/H), in line with the paradigm that pro-B cell maturation is blocked with age (Klinman and Kline, 1997). However spermidine does not affect these aging phenotypes in either spleen or bone marrow (Figure 1C-F and S1D-H). Therefore spermidine does not significantly affect hematopoiesis in old mice.

**Figure 1.**
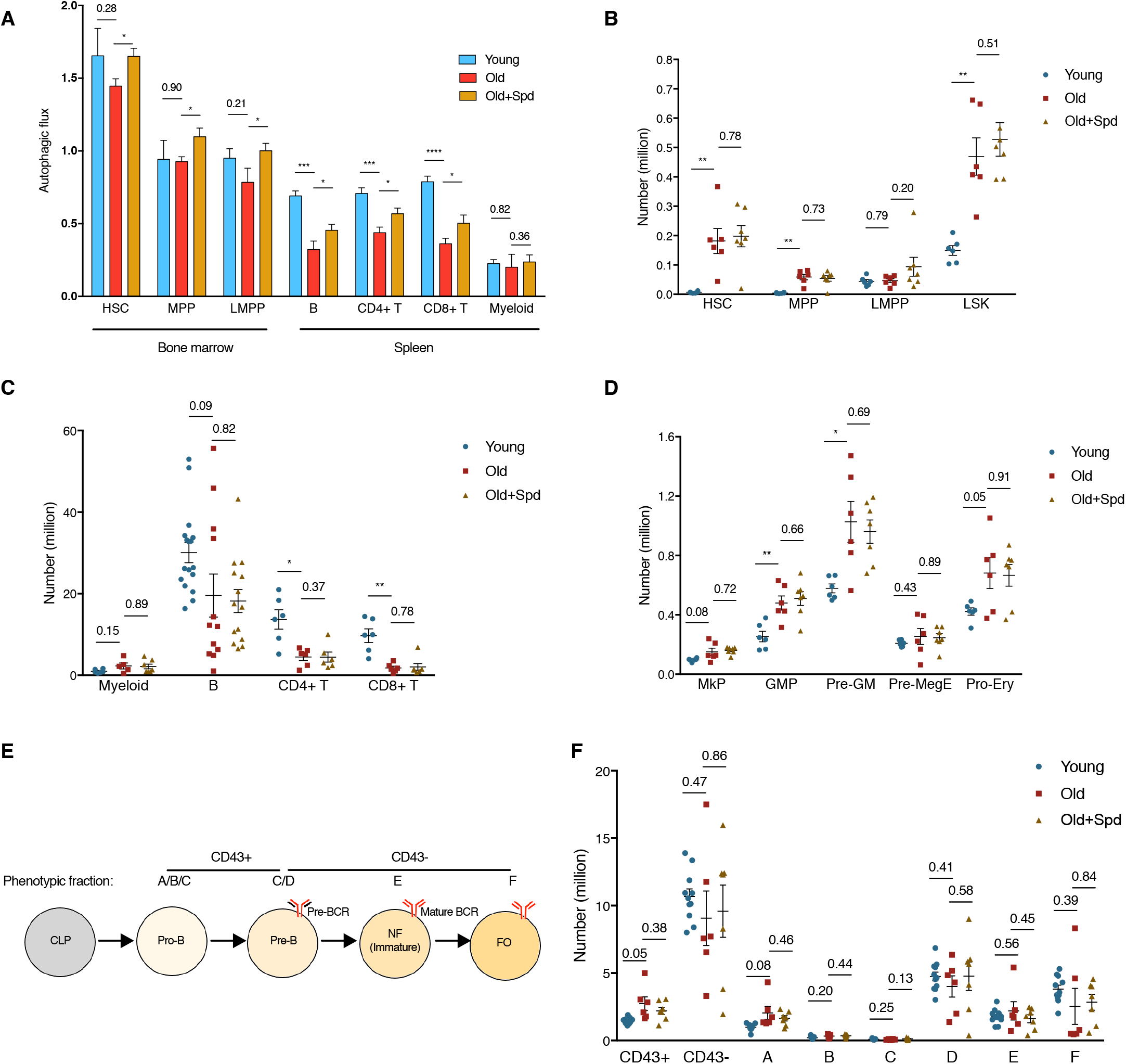
Spermidine induces autophagy *in vivo* but does not affect hematopoiesis in old mice. (A) The autophagic flux of indicated cell types from young (12 weeks) and old (22-24 months) mice, and old mice administered with spermidine for 6 weeks was measured by LC3-II staining using flow cytometry after 2 h treatment with 10 nM bafilomycin A1 (BafA1) or vehicle. Autophagic flux was calculated as LC3-II geometric mean fluorescence intensity (BafA1-Vehicle)/Vehicle. n = 5/6/7 mice for Young/Old/Old+Spd respectively. Representative histogram shown in Figure 2A. (B-F) Absolute count of indicated cell types in mice treated as in (A). (B) Expanded hematopoietic stem and progenitor cells in bone marrow (femur and tibia of both sides) of old mice. HSC, hematopoietic stem cell; MPP, multipotent progenitor; LMPP, lymphoid-biased multipotent progenitor; LSK, Lin^-^Sca1^+^cKit^+^ cell. n = 6-7 mice. (C) Old mice are lymphopenic (spleen). n = 6-17 mice as indicated by dots. (D) Expanded myeloid progenitors in bone marrow of old mice. MkP, megakaryocyte progenitor; GMP, granulocyte-macrophage progenitor; Pre-GM, pre-granulocyte/macrophage; Pre-MegE, pre-megakaryocyte/erythrocyte; Pro-Ery, pro-erythroblast cell. n = 6-7 mice. (E) Hardy fractions A-F and their correlation with B cell developmental stages. CLP, common lymphoid progenitors; Pro-B, progenitor B cells; Pre-B, precursor B cells; NF, newly formed B cells; FO, follicular B cells (mature recirculating B cells). (F) B cell development is mildly blocked at the pro-B cell stage in old mice. n = 7-11 mice, each dot represents one mouse. Data represented as mean ± SEM. Two-tailed Student’s t-test for Young/Old comparison, one-tailed Student’s t-test for Old/Old+Spd comparison (A). Two-tailed Welch’s t-test (B/C/D/F). *P<0.05, **P≤0.01, ***P≤0.001, ****P≤0.0001. See also Figure S1.

### Spermidine Rejuvenates B Cell Responses in Old Mice via Induction of Autophagy

Consistent with the flow cytometry staining of LC3-II (Figure 2A), reduced autophagic flux was confirmed in mature B lymphocytes by Western blot (Figure 2B) and by confocal microscopy using GFP-LC3 transgenic mice (Figure 2C). Old B lymphocytes accumulated LC3-II, which did not further increase by BafA1 treatment (Figure 2A-C). This indicates that autophagic flux is impaired in old cells at the level of the lysosome.

**Figure 2.**
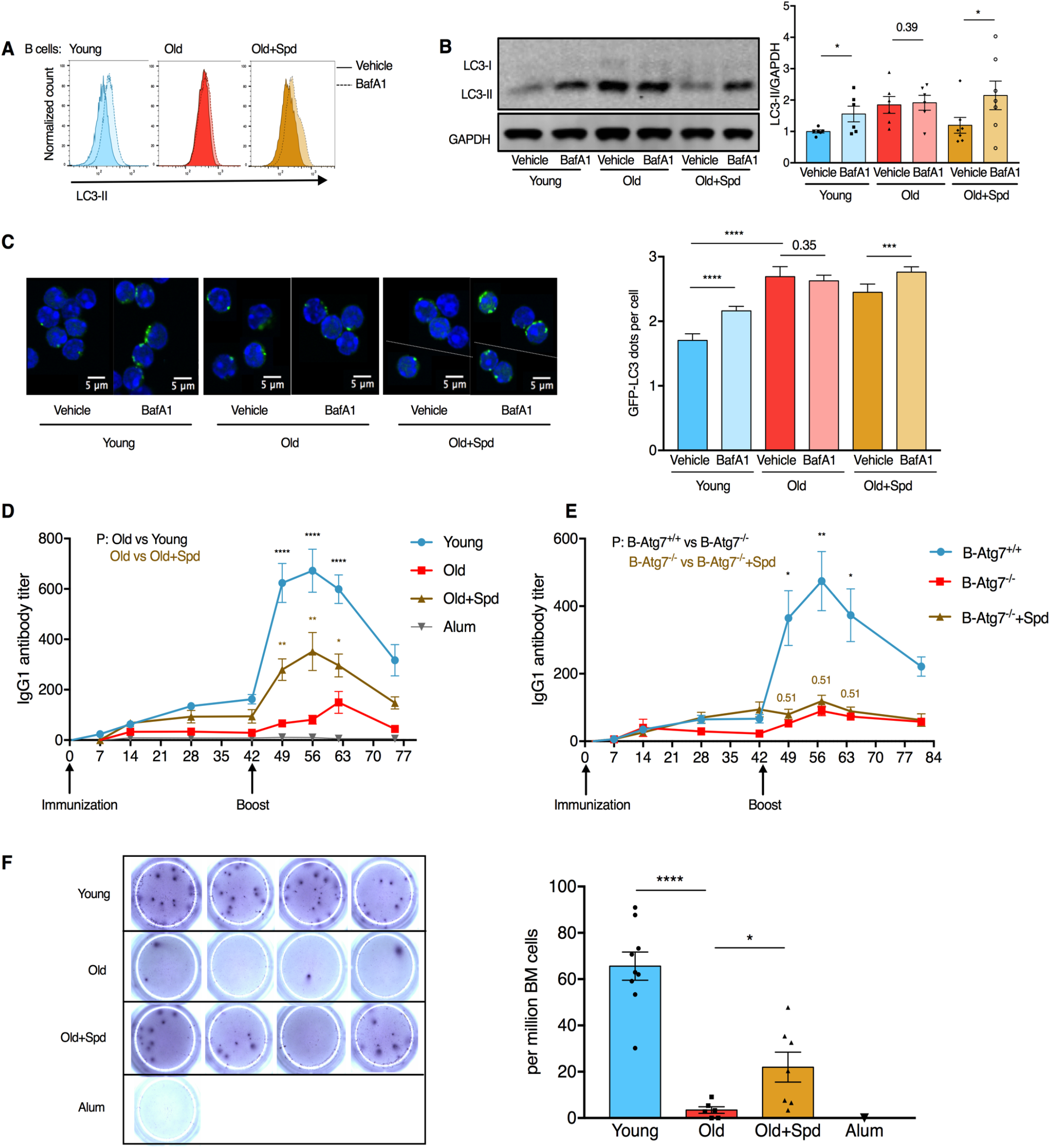
Spermidine rejuvenates B cell responses in old mice via induction of autophagy. A. Representative plot of LC3-II staining of B cells (CD19^+^).
B. LC3A/B of purified B cells from wild type mice (treated as in Figure 1A) was assessed by Western blot. The densitometric quantification is shown in the right panel, in which LC3-II was normalized to GAPDH first and then to the average of Young Vehicle. n = 6-7 mice as indicated by dots.
C. Old GFP-LC3 transgenic mice were administered with spermidine as in Figure 1A. The GFP-LC3 dots of purified B cells were measured by confocal microscopy (left) and quantified (right). N = 210-616 cells from 4-6 mice.
D. Young or old wild type mice were immunized and boosted with NP-CGG. Spermidine was given 2 days before the first immunization and then continuously in drinking water. Serum NP-specific IgG1 titers were measured by ELISA. n = 14 (Young) or 10 (Old/Old+Spd) mice from 2 experiments.
E. B cell-specific *Atg7* knockout mice (*Mb1-Cre, Atg7^f/f^*) were immunized and IgG1 responses assessed as in (D). n = 7 (B-Atg7^+/+^) or n = 3 (B-Atg7^-/-^) mice.
F. Mice from (D) were culled on day 75 and long-lived bone marrow plasma cells secreting NP-specific IgG1 were measured by ELISpot. Representative images (left) and the quantification (right) are shown. n = 6-9 mice as indicated by dots. Paired one-tailed Student’s t-test (B). Mann-Whitney test (C). Welch’s t-test (D/E/F). P values were adjusted by the Holm-Sidak method for multiple comparisons of the three time points in (D/E). Data represented as mean ± SEM. *P≤0.05, **P≤0.01, ***P≤0.001, ****P≤0.0001. See also Figure S2.

We then examined whether spermidine improves B cell function. In mice over 22 months of age, the antibody response to immunization with the model antigen NP-CGG was markedly reduced as expected, but administration of spermidine in drinking water significantly improved IgG1 responses (Figure 2D) in an autophagy-dependent manner using B cell specific conditional deletion of *Atg7* (Figure 2E, S2A/B for control for efficient deletion of *Atg7* and autophagy in B cells). At the time of sacrifice, we found very low numbers of long-lived bone marrow NP-specific plasma cells in old mice, which were significantly restored with spermidine treatment (Figure 2F). In contrast, spermidine did not induce autophagy or improve the antibody responses in young mice (Figure S2C/D). Thus, spermidine restored autophagic flux in old B lymphocytes *in vivo* and rejuvenated B cell responses in an autophagy-dependent manner.

### Spermidine Maintains Cellular Autophagy by Hypusinating elF5A

We next investigated how spermidine regulates autophagy. First, we confirmed 100 μM spermidine to be the optimal concentration to induce autophagy in the mammalian lymphocytic Jurkat cell line (Figure S3A/B) (Eisenberg et al., 2009; Morselli et al., 2011). Several signaling pathways have been described downstream of spermidine, including inhibition of histone acetyltransferases (HATs) (Madeo et al., 2018). Therefore, we investigated whether spermidine inhibits HAT activity and thereby induces *Atg7* mRNA expression as shown in yeast (Eisenberg et al., 2009). In neither of three approaches (HAT colorimetric assay, *Atg7* qPCR, and acetylated H3 pull down) did we find that spermidine had this effect (Figure S3C-E). The acetyltransferase p300 has been shown to directly acetylate certain autophagy proteins such as ATG7 to inhibit autophagy (Lee and Finkel, 2009), and spermidine was reported to inhibit p300 activity (Pietrocola et al., 2015). However, ATG7 acetylation was not affected by two tested concentrations of spermidine within 6 h of treatment (Figure S3E/F). Spermidine also failed to affect the activity of AMPK as assessed by AMPK phosphorylation (Figure S3G). However, we found that spermidine induces ER stress at the autophagy-inducing dose of 100 μM or higher (Figure S3H), demonstrated by increased levels of ATF4 (Figure S3I), phosphorylated eIF2α (Figure S3J), and increased expression of *CHOP* mRNA (Figure S3K) in Jurkat cells. Furthermore, high-dose spermidine induced cell death after 24 h (Figure S3L), presumably because spermidine levels are high in transformed cell lines and further uptake is toxic and non-specifically induces cellular stress responses, likely also inducing autophagy. ER stress was also induced by spermidine in NIH 3T3 cells as assessed by increased ATF4 expression (Figure S3M), indicating that high-dose spermidine induces cellular stress across different cell types.

Therefore in subsequent experiments, rather than adding spermidine, we opted for the depletion of spermidine either by genetic knock down of the key spermidine synthesizing enzyme ornithine decarboxylase (ODC), or with the ODC inhibitor difluoromethylornithine (DFMO) in NIH 3T3 cells (Figure 3A). Indeed, genetic knock down of *Odc* with siRNA, or DFMO treatment, reduces spermidine levels substantially, whilst supplementing cells with spermidine rescues its levels partially, as measured by GC-MS in cell lysates (Figure 3B and S4A). This demonstrates firstly that cultured cells actively synthesize spermidine via ODC, and secondly that exogenous spermidine is efficiently taken up to rescue its intracellular levels. Next, we examined pathways downstream of spermidine that had not been previously linked to autophagy. In eukaryotic cells spermidine is a unique substrate for the hypusination of the translation factor eIF5A (Figure 3A) (Rossi et al., 2014). To date, eIF5A is the only known protein containing the unusual amino acid hypusine (Dever et al., 2014). Upon knockdown of *Odc* in NIH 3T3 cells, or addition of the ODC inhibitor DFMO, we found reduced LC3-II levels and reduced eIF5A hypusination, while supplementation with spermidine rescues both (Figure 3C and S4B/C).

**Figure 3.**
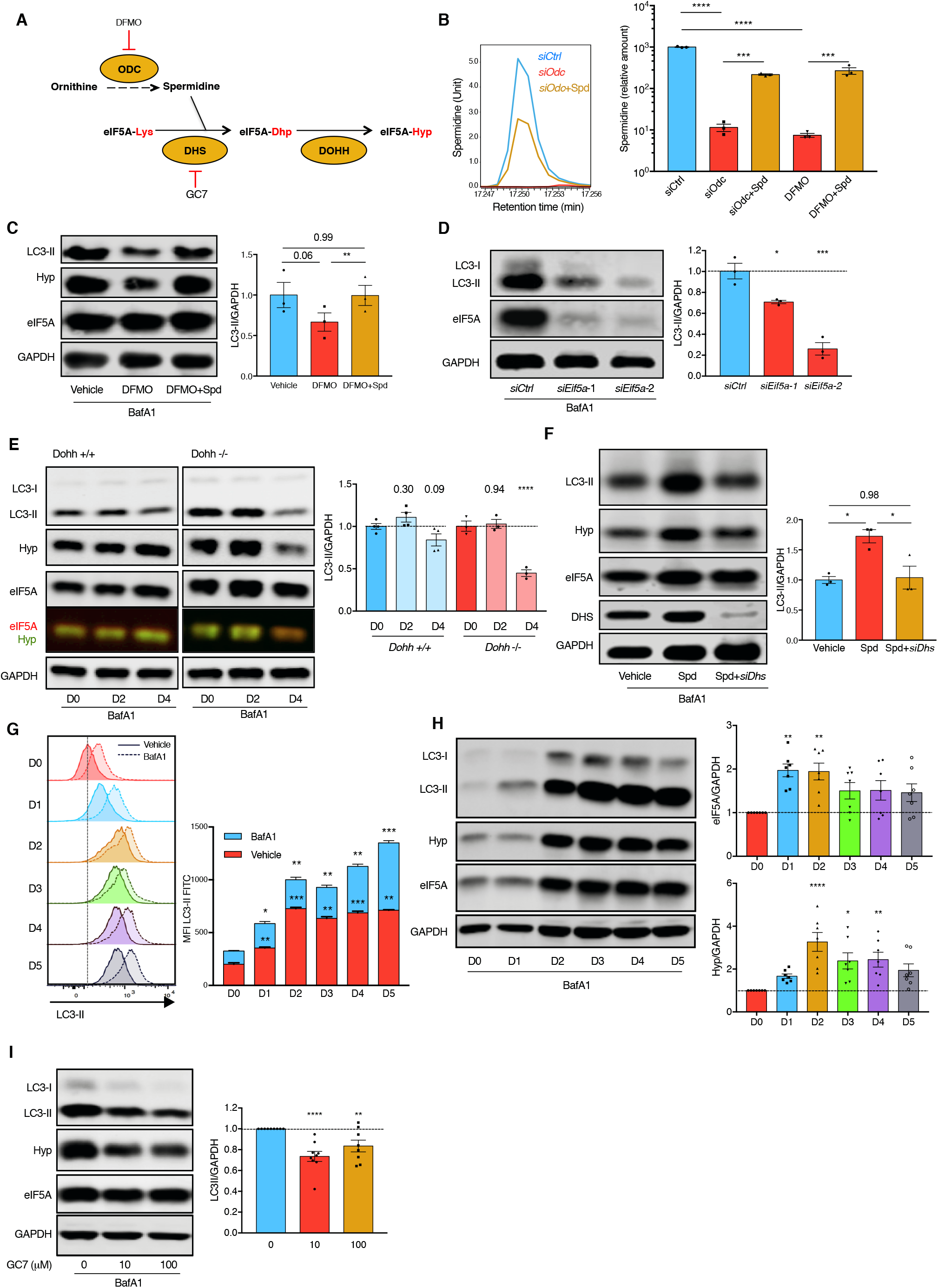
Spermidine Maintains Cellular Autophagy by Hypusinating eIF5A. (A) Spermidine synthesis and eIF5A hypusination pathway in eukaryotes. Difluoromethylornithine (DFMO) is an irreversible inhibitor of the key spermidine synthesis enzyme ornithine decarboxylase (ODC). N1-Guanyl-1,7-Diaminoheptane (GC7) is a competitive inhibitor of the key hypusination enzyme deoxyhypusine synthase (DHS). (B) NIH 3T3 cells were transfected with non-targeting control siRNA *(siCtrl)*, siRNA targeting *Odc* mRNA *(siOdc)* with/without 10 *μ*M spermidine for 3 days, or treated with 1 mM DFMO with/without 10 *μ*M spermidine for 24 h where indicated. Cellular spermidine levels were measured by gas chromatography-mass spectrometry (GC-MS) (left) and normalized to total proteins first and then to the average of *siCtrl* (right). n = 3. (C) NIH 3T3 cells were treated with DFMO in the presence/absence of spermidine for 24 h. Autophagy (LC3-II) was measured by Western blot. n = 3. (D) NIH 3T3 cells were transfected with *siCtrl* or siRNA targeting two different regions of *Eif5a* mRNA *(siEif5a-1/2)* for 3 days. LC3-II was measured by Western blot. n = 3. *siEif5a-2* was used in all other figures unless specified otherwise. (E) The knockout of *Dohh* was induced by 100 nM 4-OHT in immortalized transgenic mouse embryonic fibroblasts. LC3-II on indicated days post induction was measured by Western blot. n = 3-4. (F) Spermidine-depleted NIH 3T3 cells by *siOdc* transfection were rescued with spermidine alone or spermidine together with siRNA targeting *Dhs* mRNA *(siDhs).* LC3-II was measured 3 days post transfection by Western blot. n = 3. (G/H) B cells purified from young adult wild type mice were cultured with 10 *μ*g/mL LPS for indicated days and LC3-II was measured by flow cytometry (G) and Western blot (H). (G) The blue bars refer to BafA1 minus Vehicle. n = 3 mice. (H) The expression of overall eIF5A and hypusinated eIF5A were assessed by Western blot. n = 7 mice. (I) B cells were cultured as in G for 3 days with indicated concentrations of GC7 added on day 2 for 24 h. LC3-II was measured by Western blot. n = 6-10 mice. To measure autophagic flux, cells were treated with 10 nM BafA1 for 2 h before harvesting. Data represented as mean ± SEM. One-way ANOVA with post hoc Dunnett’s test comparing with *siCtrl* (D), D0 (G where LC3-II levels under either basal or BafA1 conditions are compared/H) or 0 *μ*M GC7 (I). One-way ANOVA with post hoc Tukey’s test (B/C/F). Two-way ANOVA with post hoc Sidak’s test (E). *P≤0.05, **P≤0.01, ***P≤0.001, ****P≤0.0001. See also Figure S3 and Figure S4.

Similarly, reduced LC3-II levels were observed when *Eif5a was* knocked down (Figure 3D and S4D-F). We next depleted the two hypusinating enzymes deoxyhypusine synthase (DHS) and deoxyhypusine hydroxylase (DOHH) to study their effect on autophagy. The knockdown of *Dhs*, knockout of *Dohh* (Pallmann et al., 2015), or GC7, a specific inhibitor of DHS reduced eIF5A hypusination and LC3-II levels (Figure 3E and S4G/H). Whilst exogenous spermidine restored LC3-II levels in cells depleted of spermidine, inhibition of eIF5A hypusination with *siDhs* or GC7 abrogated this effect (Figure 3F and S4I). Taken together the data indicates that spermidine maintains cellular basal autophagy by hypusinating eIF5A.

In activated T cells, eIF5A is one of the twenty most abundant proteins (Hukelmann et al., 2016). However, little is known about its role in lymphocytes. We measured eIF5A levels and autophagy in primary B cells during activation. LC3-II levels increased substantially over time upon activation in B cells, as measured by flow cytometry (Figure 3G) and Western blot (Figure 3H and S4J). This time-dependent increase correlated with increasing levels of both total and hypusinated eIF5A (Figure 3H). In line with our data in NIH 3T3 cells, LC3-II levels decreased with increasing doses of GC7 (Figure 3I). This pathway operates independently of mTOR whose activation inhibits autophagy. Indeed GC7 did not activate mTOR as demonstrated by S6 phosphorylation, a downstream substrate of mTOR (Figure S4K). Overall this data indicates that the eIF5A-autophagy pathway is induced upon activation of primary B cells, and physiological levels of hypusinated eIF5A are required for efficient autophagy.

### eIF5A Hypusination is Required for TFEB Expression

We next addressed how eIF5A regulates autophagy by identifying changes in expression of proteins involved in the autophagy pathway upon inhibition of eIF5A. We performed label-free quantitative mass spectrometry (MS) on nuclear and cytoplasmic fractions of activated primary B cells treated with GC7 (Figure 4A and S5A). This was followed by stable isotope labeling with amino acids in cell culture (SILAC) on activated primary B cells (Figure 4B and S5B). It has recently been reported that eIF5A regulates ATG3 protein synthesis in cell lines (Lubas et al., 2018). However ATG3 protein levels were not found reduced in primary B cells treated with GC7 as shown in our MS results (Figure 4A), and confirmed by Western blot (Figure S5C). Of the autophagy related proteins detected, TFEB was the only one repeatedly found decreased upon GC7 treatment in both approaches. TFEB is a key transcription factor regulating autophagosomal and lysosomal biogenesis (Sardiello et al., 2009; Settembre et al., 2011). The effect of inhibition of eIF5A hypusination by GC7 on TFEB was confirmed by Western blot in activated primary B cells (Figure 4C). Moreover, cellular fractionation demonstrated that TFEB protein is also decreased in the nuclear fraction (Figure 4D). Accordingly, gene expression of several TFEB targets was markedly decreased in activated B cells upon GC7 treatment (Figure 4E). Genetic knockdown of *Tfeb* also reduced autophagic flux in NIH 3T3 cells (Figure 4F and S5D) as reported (Settembre et al., 2011). Knockdown of *Eif5a* and *Dhs*, or GC7 treatment in NIH 3T3 cells led to a decrease in TFEB protein (Figure 4G/H). To test if TFEB is affected by spermidine depletion, we inhibited spermidine synthesis in NIH 3T3 cells by *siOdc* or DFMO. This caused a reduction in TFEB, which was rescued by exogenous spermidine administration (Figure 4I and S5E), suggesting that spermidine controls cellular TFEB levels. However, spermidine failed to maintain TFEB levels when *Eif5a* was knocked down or eIF5A Figure 4. eIF5A Hypusination is Required for TFEB Expression. hypusination was inhibited (Figure 4J and S5F/G), indicating that the regulation of TFEB by spermidine is mediated via eIF5A hypusination.

**Figure 4.**
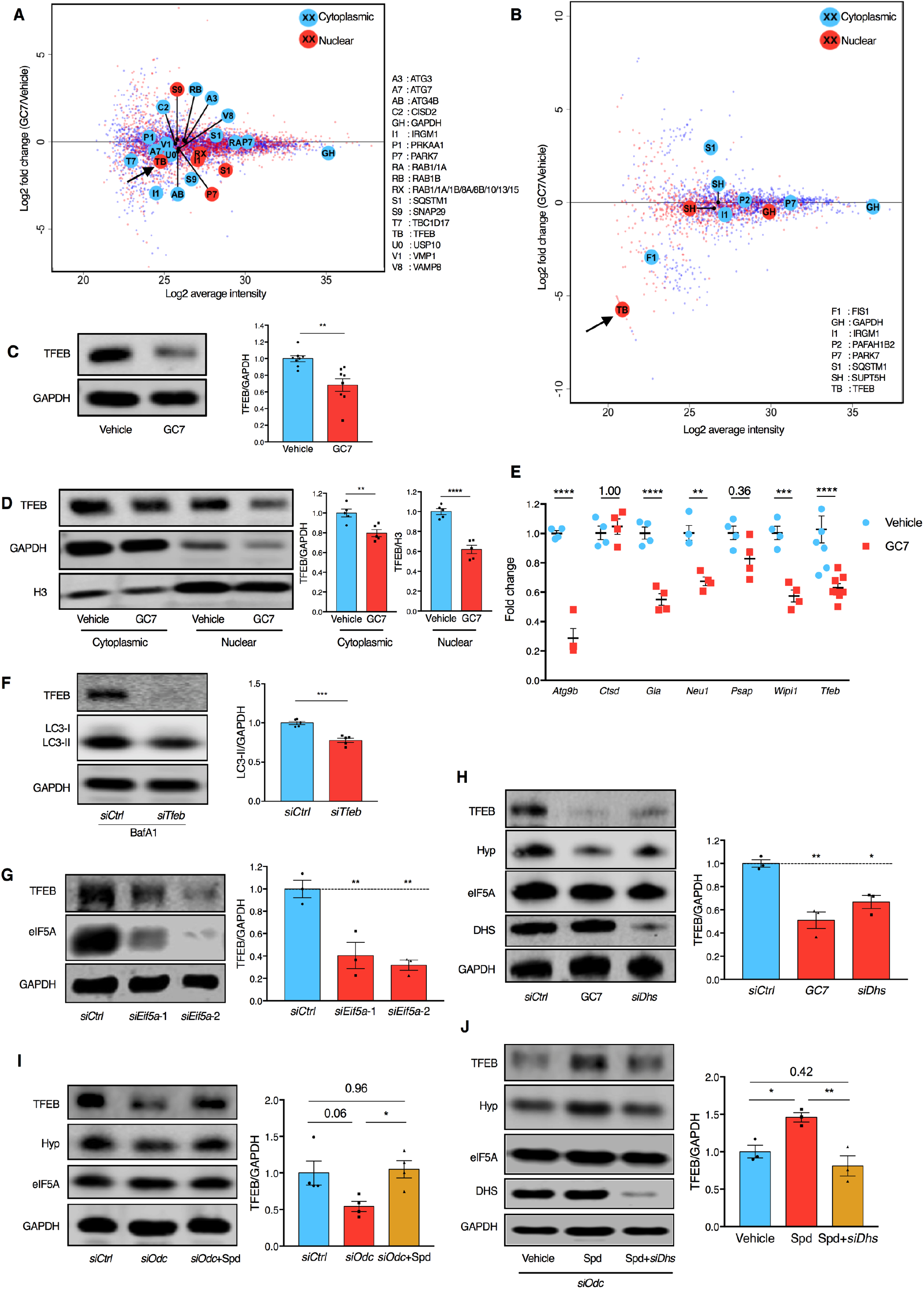
eIF5A Hypusination is Required for TFEB Expression. (A) Murine B cells were treated with 10 *μ*M GC7 for 24 h as in Figure 3I and fractionated for label-free quantitative protein mass spectrometry (MS) analysis. Identified autophagy proteins are highlighted by enlarged, annotated circles, and TFEB (TB) is further indicated by a black arrow. (B) Murine B cells were cultured in medium containing amino acids with heavy isotope labeling (GC7-treated) or light isotopes (Vehicle). Cells that had divided four times were sorted by flow cytometry and mixed at 1:1 ratio (heavy: light isotope labelling), followed by cell fractionation and protein MS analysis. For the repeat the labeling was swapped. Data represent the average of protein changes from the two repeats. Identified autophagy proteins are highlighted as in A. (C-E) Murine B cells were treated with GC7 as in A. The overall TFEB (C), and cytoplasmic/nuclear TFEB (D) were assessed by Western blot. n = 3 mice. (E) The expression of TFEB-target genes was measured by qPCR with *Gapdh* as reference gene. n = 4-8 mice as indicated by dots. (F) NIH 3T3 cells were transfected with *siCtrl* or siRNA targeting *Tfeb* mRNA *(siTfeb)* for 3 days. BafA1 was added 2 h before harvest. LC3-II was measured by Western blot. n = 5. (G) NIH 3T3 cells were transfected with *siEif5a* for 3 days. TFEB expression was measured by Western blot. n = 3. (H) NIH 3T3 cells were transfected with *siDhs* for 3 days or treated with 100 *μ*M GC7 for 24 h. TFEB expression was measured by Western blot. n = 3. (I) NIH 3T3 cells were transfected with *siOdc*, and treated with or without 10 *μ*M spermidine for 3 days. TFEB expression was measured by Western blot. n = 4. (J) NIH 3T3 cells were transfected with *siOdc* to deplete endogenous spermidine and treated with 10 *μ*M spermidine alone or in combination with *siDhs* transfection. TFEB expression was analyzed by Western blot. n = 3. Data represented as mean ± SEM. Student’s t-test (C/F). Two-way ANOVA with post hoc Sidak’s test (E). One-way ANOVA with post hoc Dunnett’s test comparing with *siCtrl* (G/H), or with post hoc Tukey’s test (I/J). *P≤0.05, **P≤0.01, ***P≤0.001, ****P≤0.0001. See also Figure S5.

### Hypusinated eIF5A Regulates TFEB Synthesis

TFEB is a very short-lived protein (< 4 hours half-life) (Xiao et al., 2015) compared to other proteins in the cell (median 36 hours half-life) (Cambridge et al., 2011), and is therefore expected to require more active translation than most proteins. We first investigated if eIF5A controls overall protein synthesis in lymphocytes as previously reported (Landau et al., 2010). We measured translation by flow cytometry and found that inhibition of eIF5A hypusination by GC7 caused a reduction in protein synthesis rate in activated primary B cells (Figure 5A). Protein levels reflect the balance between protein synthesis and degradation. We therefore measured protein synthesis regulated by eIF5A hypusination alone using ribosome profiling in activated primary B cells, the first study of this kind in mammalian cells (Schuller et al., 2017; Woolstenhulme et al., 2015). We found that GC7 treatment led to stalled ribosomes at both initiation and termination sites (Figure 5B/C). The stalled translation initiation may be a result of increased eIF2α phosphorylation and reduced 4E-BP phosphorylation caused by GC7 as reported (Landau et al., 2010). Stalling of ribosomes at the triproline motif (PPP) was found in activated primary B cells (Figure 5D), as observed in yeast and bacteria (Doerfel et al., 2013; Schuller et al., 2017; Ude et al., 2013), indicating that polyproline is a conserved ribosome-pausing motif across kingdoms. Interestingly both human and mouse TFEB have at least one triproline motif (Figure 5E). Indeed sequences around this triproline are also ribosome-pausing motifs (SPP and PPV) (Schuller et al., 2017), suggesting that this triproline motif may confer TFEB the specific requirement to rely on eIF5A for smooth translation. Therefore, we examined if the triproline Figure 5 Hypusinated eIF5A regulates TFEB synthesis. motif in murine/human TFEB is sufficient to affect translation rate as regulated by hypusinated eIF5A. We generated three different constructs, using labile mCherry expression to report on the synthesis of either the TFEB triproline-containing motif, 13 consecutive prolines (13Pro) or a random sequence (Figure 5E). After transfection into NIH 3T3 cells, we measured translation using the ratio of the reporter mCherry to the transfection control GFP. As expected, the ratio of mCherry/GFP was mildly reduced by GC7 for the random sequence and reduced by nearly half for the 13Pro motif, while the synthesis of TFEB triproline motif was also significantly inhibited (Figure 5F), indicating that the TFEB triproline motif requires hypusinated eIF5A for efficient synthesis. Moreover, mutating the triproline motif (PPP to AAA) partially rescued the expression of TFEB in the absence of hypusinated eIF5A (Figure 5G). These data suggests that hypusinated eIF5A directly facilitates the synthesis of the TFEB triproline-containing motif.

**Figure 5.**
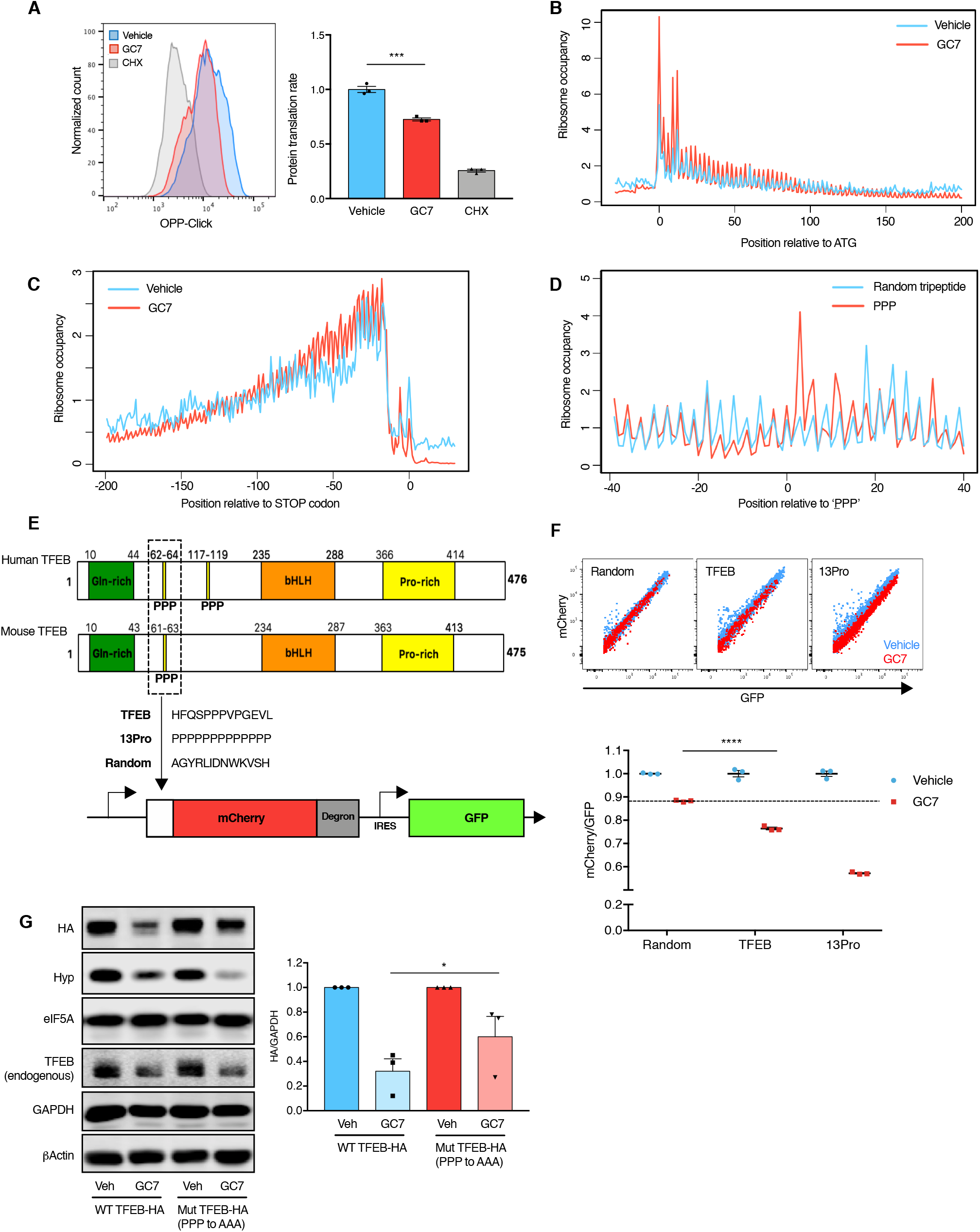
Hypusinated eIF5A Regulates TFEB Synthesis. (A) LPS-stimulated murine splenocytes were treated with 10 *μ*M GC7 for 24 h as in Figure 3I or with the protein synthesis inhibitor cycloheximide (CHX) as control for 2 h. The relative protein synthesis rate of B cells (B220^+^CD19^+^) was measured by OPP-Click assay with flow cytometry and normalized to Vehicle. n = 3. (B/C) Murine B cells were treated with 10 *μ*M GC7 as in Figure 3I and processed for ribosome profiling. Ribosome occupancy of all genes is aligned at the start codon (B) or stop codon (C) and normalized to a mean value of 1 for each gene. (D) Murine B cells were cultured with LPS for 3 days and processed for ribosome profiling. Ribosome occupancy of all genes expressing PPP motifs is aligned at the underlined Pro on the P site of the ribosome and normalized to a mean value of 1 for each gene. (E) The polyproline motif of TFEB with its surrounding sequence was inserted before mCherry-degron to report on protein translation. GFP after IRES is used to report on transfection. 13 consecutive prolines (13Pro) was used as positive control for translational stalling, while 13 random amino acids (Random) was used as negative control. (F) NIH 3T3 cells were transfected with the plasmids in (E) for 24 h, treated together with GC7 or Vehicle. The expression of GFP and mCherry was measured by flow cytometry (upper panel). The ratio of geometric mean intensity mCherry/GFP was normalized to the average of Vehicle (lower panel). n = 3. (G) NIH 3T3 cells were transfected with the plasmids expressing wild type (WT TFEB-HA) or mutant (Mut TFEB-HA, PPP to AAA) murine TFEB fused with an HA tag at C-terminus for 24 h, treated together with GC7 or Vehicle. The expression of HA-tagged TFEB was assessed by Western blot (left) and quantified by normalizing HA to GAPDH (right). n = 3. Data represented as mean ± SEM. Student’s t-test (A/F) or ratio paired t-test (G). *P≤0.05, ***P≤0.001, ****P≤0.0001.

### Hypusination of eIF5A is Essential for B Cell Development and Activation

eIF5A and its hypusination are essential for cellular growth in cultured cells (Park et al., 1994) and early embryonic development in mice (Nishimura et al., 2012), but its tissue-specific function is still unclear, although inducible whole-body knockout of *Dhs* in mice leads to a fatal wasting syndrome (Pallmann et al., 2015). To investigate the function of hypusinated eIF5A during B cell development and activation *in vivo*, we generated competitive mixed bone marrow chimeras with inducible deletion of the hypusinating enzyme DHS. *Dhs* deletion was induced after long-term engraftment and B cell lineage reconstitution of CD45.2^+^ cells was examined on day 8 and 30 after deletion (Figure 6A and S6A/B). Percentages of CD45.2^+^ of both transitional and mature follicular circulating B cells in peripheral blood were significantly affected by *Dhs* deletion on day 30 (Figure 6B). Upon sacrifice of the mice on day 34 we investigated if this was due to a loss of progenitors in the bone marrow. Unexpectedly, all multipotent hematopoietic progenitors investigated (HSC, MPP, LMPP, LSK) were severely affected by DHS depletion (Figure 6C). This is in line with a bone marrow hypocellularity reported previously in whole body *Dhs* knockout mice (Pallmann et al., 2015). Consistent with depleted HSCs, splenic CD45.2^+^ myeloid cells, which mostly lack self-renewal capacity and rely on replenishment from progenitor cells, were also severely depleted after *Dhs* deletion (Figure S6C). Similarly, early B cell progenitors (pre-pro B cells), newly formed B cells and mature B cells from bone marrow as well as splenic transitional, marginal zone and follicular B cells were found significantly diminished (Figure 6D/E). We checked if the remaining CD45.2^+^ cells were indeed deleted for *Dhs* in blood (Figure S6D), bone marrow and spleen (Figure S6E). This analysis revealed that whilst on day 8 after deletion with Tamoxifen, blood cells were adequately deleted in the floxed *Dhs* mice, in cells that survived to day 34, *Dhs* was not deleted in most mice (Figure S6E). The overall cellularity in bone marrow and spleen was not affected despite of the reduced CD45.2^+^ cell numbers (Figure S6F), indicating that the remaining CD45.1^+^ and non-deleted CD45.2^+^ cells expanded and filled up the niche. The data together demonstrates a profound requirement of the eIF5A pathway in hematopoiesis and long-term survival of mature B cells.

To circumvent the defects of hematopoiesis and assess the role of eIF5A hypusination in B cell activation, we further attempted to delete *Dhs* after the first immunization with NP-CGG in *Dhs* KO bone marrow chimeric mice supported with Rag^-/-^ bone marrow cells (to provide wild type hematopoietic cells for the survival of the mice, without T and B cells). As before, after 3 weeks of deletion, all remaining lymphocytes expressed wild type *Dhs* only. Levels of NP-specific IgG1 post-boost were not significantly changed in KO mice (data not shown), which can be attributed to contamination from non-deleted memory B cells. Overall the data implies that complete inhibition of eIF5A hypusination kills hematopoietic cells, and its effect on generation of antibodies cannot easily be investigated *in vivo* due to a) the rapid elimination of cells knocked out for *Dhs* and b) the limiting deletion efficiency in mature B cells with inducible deletion models.

**Figure 6.**
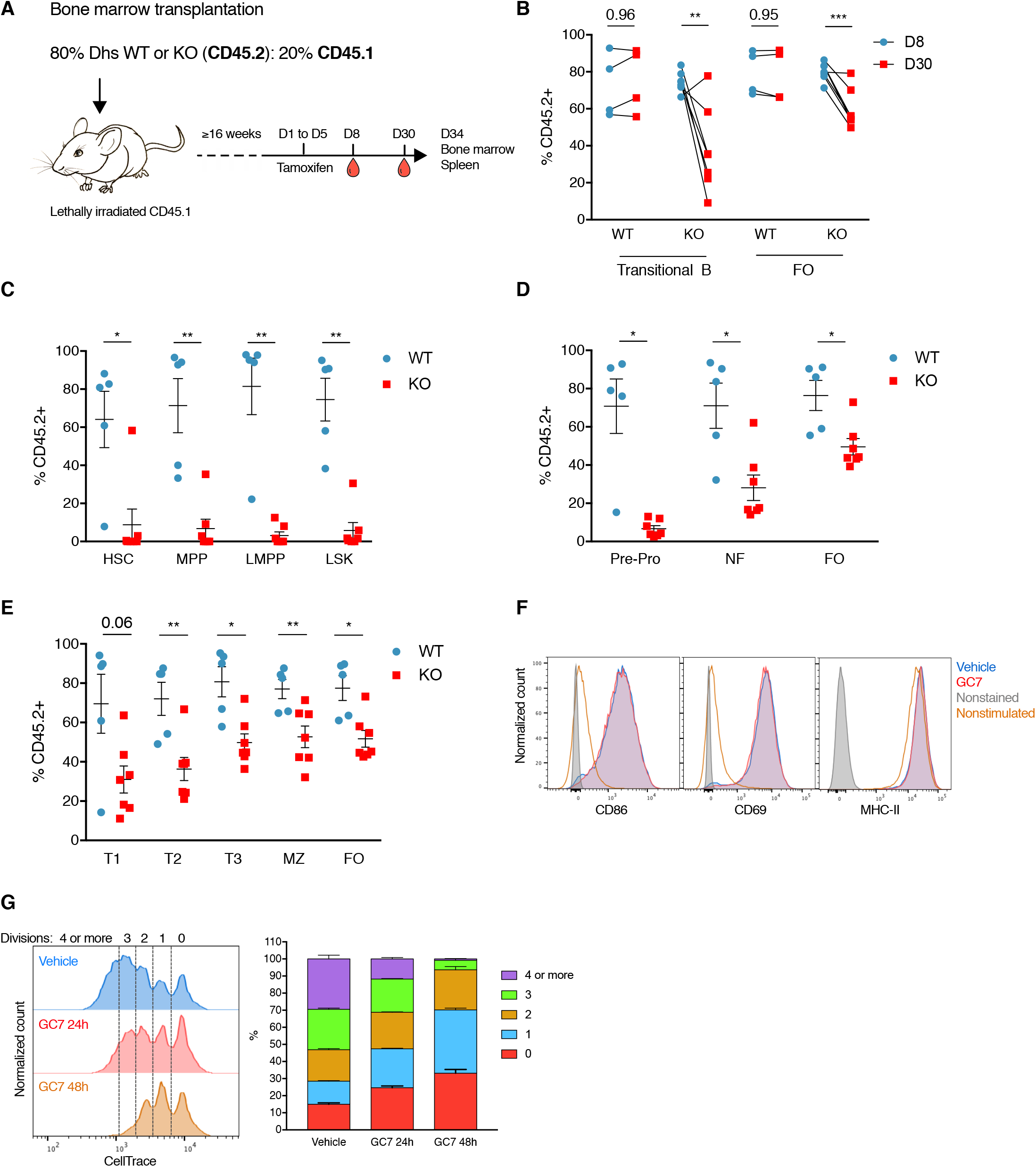
Hypusination of elF5A is essential for B cell development and activation. (A-E) Competitive bone marrow chimeric mice were generated by transplanting bone marrow cells from tamoxifen-inducible CD45.2^+^ *Dhs* knockout mice (WT: CAG-Cre/Esr1^+^, *Dhs*^+I+^; KO: CAG-Cre/Esr1^+^, *Dhs*^f/f^,) and wild type CD45.1^+^ competitors into wild type CD45.1^+^ recipient mice (A). After ≥ 16 weeks of long-term reconstitution, tamoxifen was administered by oral gavage for 5 consecutive days, followed with lineage contribution assessment in peripheral blood (B), bone marrow (C/D), and spleen (E). (B) The contribution of CD45.2^+^ cells to transitional (CD19^+^CD93^+^) and mature follicular (FO) B cells (CD19^+^CD93^-^) in peripheral blood on day 8 and day 30 post tamoxifen induction was assessed by flow cytometry. n = 4-7 mice as indicated by dots. Two-way ANOVA with post hoc Sidak’s test for transitional B cells and follicular B cells separately. (C-E) The contribution of CD45.2^+^ cells to bone marrow hematopoietic stem and progenitor cells (C), pre-pro B cells (Hardy fraction A), newly formed B cells (NF, fraction E), follicular B cells (FO, fraction F) (D), and spleen transiational B cells (T1-3), marginal zone B cells (MZ) and follicular B cells (E) was assessed by flow cytometry on day 34 post tamoxifen induction. n = 5-7 mice as indicated by dots. Welch’s t-test. *P≤0.05, **P≤0.01, ***P≤0.001. (F) Wild type murine B cells were stimulated with LPS for 1 day with or without 10 *μ*M GC7. The expression of early activation markers CD86, CD69, and MHC II was assessed by flow cytometry. (G) Wild type murine B cells were stained with CellTrace and cultured with LPS for 3 days. GC7 was added 24 h or 48 h before harvest for cell proliferation analysis (left). The percentage of cells that divided for the indicated times is quantified (right). n = 6 mice. Data represented as mean ± SEM. See also Figure S6.

To investigate if eIF5A has an effect on B cell activation, we induced B cell activation *ex vivo* with LPS while inhibiting hypusination with GC7. The up-regulation of early activation markers, including CD86, CD69, and MHC-II (Figure 6F and S6G) were not changed upon GC7 treatment, indicating that hypusinated eIF5A does not regulate early activation signaling pathways. However, activation-induced cell proliferation was severely impaired upon inhibition of hypusinated eIF5A (Figure 6G) in a time-dependent fashion, in line with the known function of eIF5A in supporting cellular growth (Park et al., 1994). Taken together, these results indicate that the novel pathway identified here is not required for the signaling pathway immediately downstream of the LPS sensor TLR4, but is required for more distal signaling events leading to proliferation and other events for which autophagy might be required such as maintenance of organelle homeostasis and persistent antibody production (Pengo et al., 2013).

### Spermidine Induces TFEB Expression and Improves the Function of Old Human B Cells

To investigate if this pathway regulates aging in human B cells, we firstly measured the levels of hypusinated eIF5A and TFEB in PBMCs from healthy donors of different ages. We found that PBMCs from donors aged 65 and over show a distinct reduction of TFEB protein. In some aged donors TFEB was even undetectable by Western blot (Figure 7A). In contrast, the mRNA of *TFEB* was not reduced in PBMCs from old donors (Figure 7B), further indicating that the diminished TFEB protein expression in the elderly is due to translational defects. Not only hypusinated eIF5A, but also overall eIF5A was significantly diminished in human PBMCs from donors ≥65 (Figure 7A), suggesting that cells may coordinate the expression of overall eIF5A with their hypusination levels *in vivo*. Indeed TFEB protein levels correlate well with the expression of eIF5A (Figure S7A). Using mass spectrometry, we show that endogenous spermidine in human PBMCs declines with age (Figure S7B) as reported (Pucciarelli et al., 2012). Next, we investigated whether TFEB is regulated by spermidine and eIF5A in human primary B cells. Human B cells from young donors stimulated with anti-IgM and CD40L and treated with DFMO *ex vivo* showed reduced eIF5A hypusination and TFEB, which were rescued by exogenous spermidine supplementation (Figure 7C).

**Figure 7.**
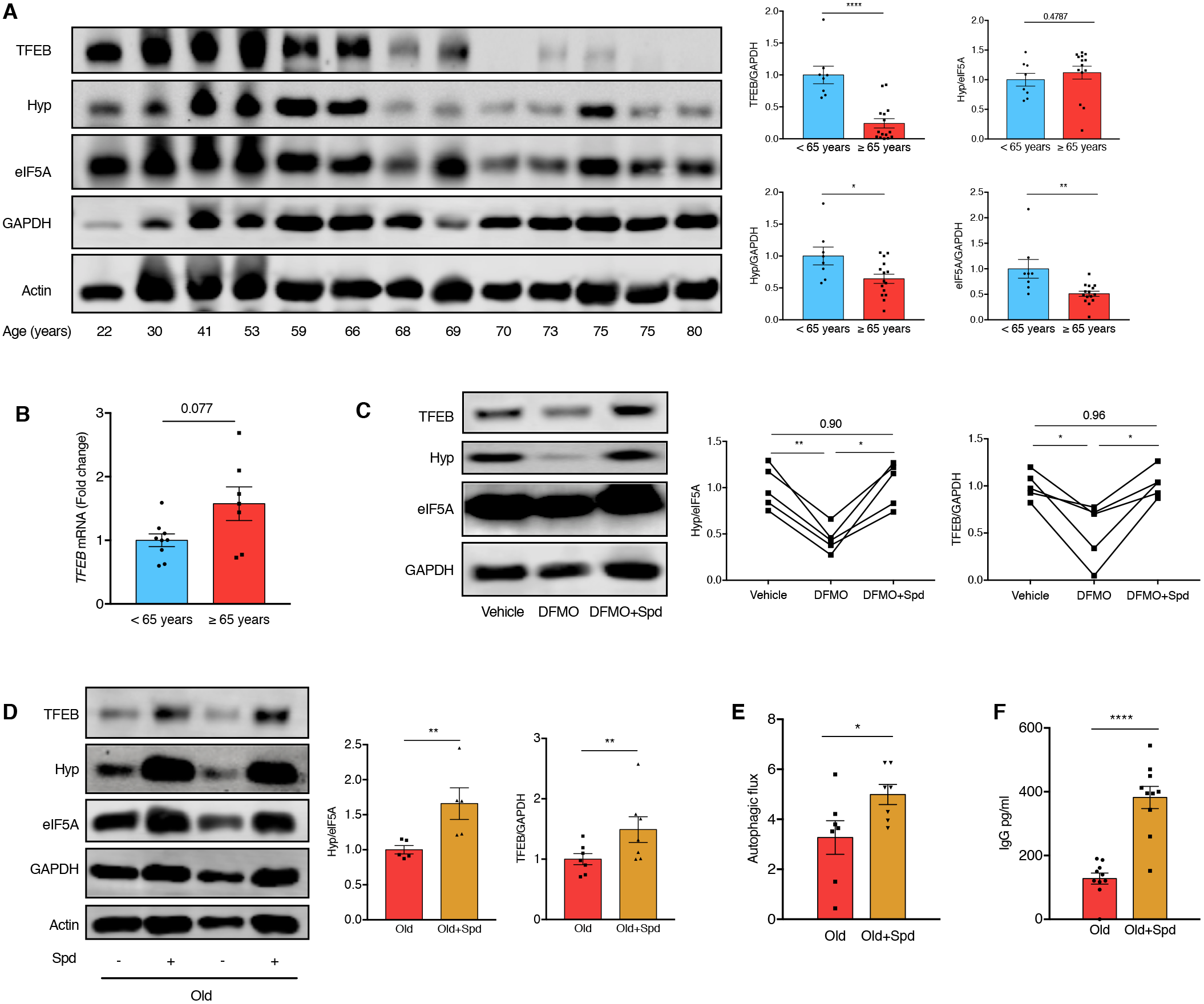
Spermidine induces TFEB expression and improves the function of old human B cells. (A) The protein levels of TFEB, hypusinated eIF5A, and total eIF5A of PBMCs from healthy human donors of indicated ages were assessed by Western blot. A representative plot (left) and quantifications (right) are shown. Target band intensity was normalized to GAPDH or eIF5A first and then to the average of the < 65 years group. n = 8 (<65 years) or 15 (≥65 years) donors as indicated by dots. Student’s t-test. (B) *TFEB* mRNA of human PBMCs was measured by qPCR with *GAPDH* as reference gene. n = 9 (<65 years) or 7 (≥65 years) donors as indicated by dots. Welch’s t-test. (C) Sorted B cells from young human donors were cultured with anti-IgM and CD40L for 7 days treated with DFMO alone or together with 10 *μ*M spermidine. The levels of TFEB and eIF5A hypusination were assessed by Western blot (left panel) and quantified (middle/right panels). n = 5 donors as indicated by dots. Paired oneway ANOVA with post hoc Tukey’s test. (D-F) Sorted B cells from old human donors (age 77.5 ± 6.3 years) were cultured as in B together with 10 *μ*M spermidine. (D) The protein levels of TFEB and eIF5A hypusination were measured by Western blot. Target band intensity was normalized to eIF5A (for Hyp) or GAPDH (for TFEB) first and then to the average of the Old group. n = 5 (Hyp/eIF5A) or 7 (TFEB/GAPDH) donors. (E) Autophagic flux was measured by flow cytometry as in Figure 1A. n = 7 donors. (F) Supernatant IgG was assessed by ELISA. n=10 donors. Paired one-tailed (D/E) or two-tailed (F) Student’s t-test. Data represented as mean ± SEM. *P≤0.05, **P≤0.01, ***P≤0.001, ****P≤0.0001. See also Figure S7.

Lastly and most importantly, we tested if spermidine improves old human B cell responses. The levels of both hypusinated eIF5A and TFEB (Figure 7D), as well as autophagic flux (Figure 7E) were significantly restored by spermidine treatment in *ex vivo* cultured B cells from aged donors in which spermidine levels are naturally low. IgG production by B cells from aged donors was also improved (Figure 7F). Thus, replenishment of spermidine in B cells from aged donors rejuvenated their function.

## DISCUSSION

This study demonstrates that autophagic flux is regulated translationally. We identified the precise mechanism of this regulation, whereby the metabolite spermidine is a substrate for the hypusination of the translation factor eIF5A, which in turn controls the synthesis of TFEB protein via its triproline motif. This mechanism operates normally in young B cells, in which spermidine is abundant. In old B cells, however, replenishing spermidine levels restored this pathway, thereby rejuvenating responses and function *in vivo* in mice and *ex vivo* in human. Depleted spermidine levels and subsequent low TFEB expression may be an important cause of the reduction in autophagy in the aging adaptive immune system, as well as in other tissues. However, our study does not exclude that other potential direct or indirect mechanisms may play a role in the anti-aging effects of polyamines such as epigenetic modifications or changes in metabolism. It is possible that multiple mechanisms including autophagy induction are involved spermidine’s rejuvenation effects.

### Translation and Autophagy Coordinate with Each Other

Not much is known about translation in lymphocytes except for a few pioneering studies in T (Hukelmann et al., 2016; Tan et al., 2017) and B cells (Diaz-Munoz et al., 2015). Naïve B cells are in a metabolically quiescent state, with low protein synthesis (eIF5A expression) and autophagy, both of which are quickly up regulated after activation. Autophagy and the translation machinery may form a regulatory loop to support the metabolic requirements of proliferation and immunoglobulin production following activation. In this loop, autophagy provides substrates and energy for translation, while increased translation induces the synthesis of certain autophagic proteins such as TFEB. Specifically, a particular requirement for rapid provision of amino acids and unfolded protein degradation by autophagy may exist in long-lived plasma cells, which secrete antibodies at the rate of about 2,000 molecules per second (Clarke et al., 2015; Pengo et al., 2013).

### Translational Control of TFEB

The anti-aging function of TFEB seems to be highly conserved, found in mice and humans. Earlier studies found it to be a uniquely important transcription factor for lifespan extension in *C. elegans* (Lapierre et al., 2013). Moreover, it is likely that the control of TFEB via translation is a broad, maybe universal mechanism, operating across many tissue types (here in NIH 3T3 cells and in mouse and human primary B cells). TFEB is known to be post-translationally controlled via (1) mTORC1-dependent phosphorylation for cytoplasmic retention, and (2) calcium-dependent dephosphorylation to signal for nuclear translocation (Napolitano and Ballabio, 2016). We now add an mTOR-independent step of regulation at the level of translation. Interestingly, our data suggest that the eIF5A pathway controls synthesis of proteins containing PPP motifs, which are highly specific binding motifs with low affinity and as such a hallmark of transcription factors (Morgan and Rubenstein, 2013).

ATG3 protein translation was recently reported to be controlled by hypusinated eIF5A in cell lines (Lubas et al., 2018), However, we could not find a reduction of ATG3 with GC7 treatment in primary B cells indicating that this regulation may be cell type specific (Figure 4A and S5C). MHC II also contains polyproline motifs but did not reduce with GC7 treatment either (Figure 6F and S6G). These observations suggest that the presence of a polyproline motif alone is insufficient to cause reduced protein expression by inhibition of hypusinated eIF5A. We hypothesize that a stronger stalling motif (such as SPPPVP in TFEB), and a short half-life contribute to diminished protein expression in the absence of hypusinated eIF5A, and predict that other such autophagy proteins are regulated by eIF5A.

### Harness the Pathway to Rejuvenate Human Aging

We have shown eIF5A and TFEB levels are correlated with age in immune cells. Blood aging biomarkers are urgently needed to gauge the efficacy of new drugs that might prolong health span. Autophagy is one of the few general mechanisms that underpin many age-related diseases and is therefore a good target for anti-aging drugs. Lastly given that the translation factor eIF5A is regulated by post-translational modifications, mediated by enzymes, these can potentially be harnessed therapeutically.

## STAR METHODS

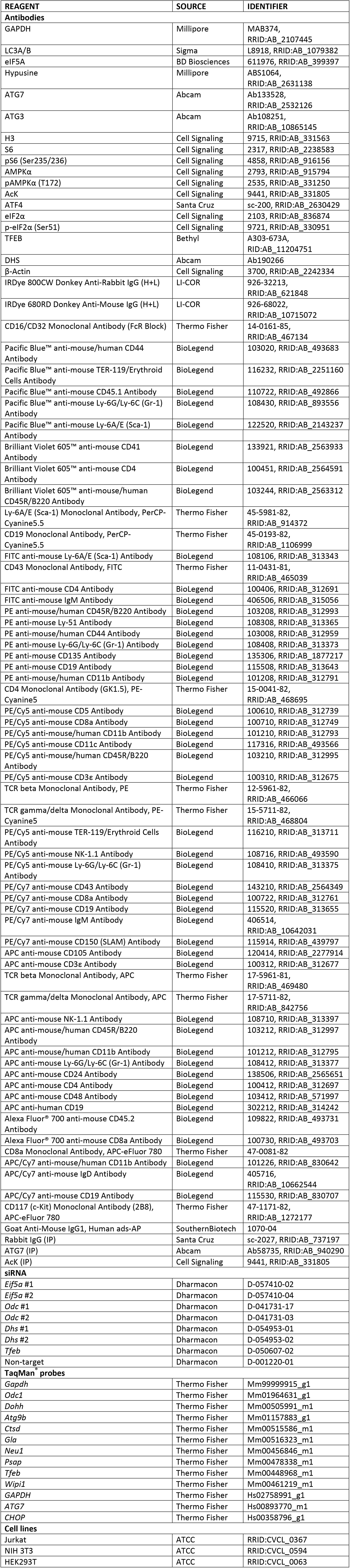
Key resource table.

### Mice

Young (6-12 weeks) and old (22-24 months) C57BL/6J wild type mice (RRID:IMSR_JAX:000664) were purchased from Charles River UK. CD45.1^+^ B6.SJL mice (RRID:IMSR_JAX:100012) were purchased from Charles River UK and bred in BSU at the Kennedy Institute of Rheumatology. *Mb1-cre* mice were a kind gift from M Reth, *Atg7^flox^* mice from M Komatsu (1year old mice were used in this study). GFP-LC3 mice from N Mizushima and Stefan Balabanov made the bone marrow from inducible *Dhs^flox^* mice available (Pallmann et al., 2015). 5 mM Spermidine (CAY14918, Cambridge Bioscience) was administered to mice in the drinking water, changed three times per week. All mice were held at Biomedical Services, University of Oxford. Animal experiments were approved by the local ethical review committee and performed under UK project licenses PPL 30/2809 and then PPL 30/3388.

### Bone marrow chimera

Bone marrow cells were collected from the femur and tibia of the donor mice (*Dhs^+/+^* or *Dhs*^-/-^ CD45.2^+^, with young B6.SJL CD45.1^+^ mice as competitors) and frozen in cryopreservation medium for long-term storage. Carefully thawed cells were counted and mixed at the indicated ratio for intravenous (iv) injection. The recipient B6.SJL mice were lethally irradiated (450 cGy twice, 4 hours apart) and rested for 1 hour prior to injection. A total of 1.5 million bone marrow cells were injected intravenously into recipient mice. After at least 16 weeks of long-term reconstitution, tamoxifen (Sigma) dissolved in corn oil was administered via oral gavage (5 mg/mouse/day for 5 consecutive days) and peripheral blood and organs were collected for analysis.

### Mouse immunization

Mice were injected intraperitoneally (ip) with 50 μg NP-CGG (N-5055D-5, Biosearch Technologies, dissolved in PBS) in Imject Alum adjuvant (Thermo Fisher) on day 0 followed by secondary immunization on or after day 42 (50 μg NP-CGG in PBS ip). Peripheral blood samples were collected from tail vein on indicated days.

### Mouse B cell purification and stimulation

RBC-lysed mouse splenocytes were purified using the Pan B Cell Isolation Kit II (130-104-443, Miltenyi). Cells were cultured at a density of 1 million/ml supplemented with 10 μg/ml LPS (Santa Cruz) for stimulation. Medium was replaced on day 3 of culture by replacing half volume with fresh culture medium containing LPS followed by analysis on day 4 and day 5.

### Western blot

Cells in suspension were washed with PBS and lysed using NP-40 lysis buffer containing proteinase inhibitors (Sigma) and phosphatase inhibitors (Sigma) on ice. After spinning down the debris, protein concentration in the supernatant was quantified by BCA Assay (23227, Thermo Fisher). Reducing Laemmli Sample Buffer was then added to the supernatant and heated at 100°C for 5 minutes. 5-20 μg proteins were loaded for SDS-PAGE analysis. For LC3 western blot, 15% Tris-HCl gel and SDS running buffer was used to separate LC3-I and LC3-II. For non-LC3 western blot, NuPAGE Novex 4-12% Bis-Tris gradient gel (Thermo Fisher) with MES running buffer (Thermo Fisher) was used. Proteins were transferred to a PVDF membrane (IPFL00010, Millipore) and blocked with 5% skimmed milk-TBST. Membranes were incubated with primary antibodies dissolved in 1% milk overnight and secondary antibodies dissolved in 1% milk with 0.01% SDS for imaging using the Odyssey CLx Imaging System. Data were analyzed using Image Studio Lite.

### Immunoprecipitation

After various treatments an equal number of cells were collected, washed with PBS and lysed using NP-40 lysis buffer with proteinase and phosphatase inhibitors on a rotator at 4°C for 20 minutes., 950 μL lysis buffer was used for 25 million Jurkat cells. Cell debris was centrifuged down at 13000 rpm for 15 minutes at 4°C. 50 μL supernatant was collected as whole cell lysate (input control), and the protein concentration was measured by BCA assay. The remaining supernatant was mixed with 15 μL Protein A Agarose beads (Thermo Fisher) in PBS, then rotated at 4°C for 15 minutes to pre-clear the lysate. The beads were removed by centrifugation at 6000 rpm for 1 minute. Immunoprecipitation (IP) antibody (AcK, ATG7, or rabbit IgG) and 20 μL Protein A agarose beads were added to the lysate and then rotated at 4°C overnight. The beads were washed twice with NP-40 lysis buffer containing proteinase and phosphatase inhibitors and the supernatant discarded. 35 μL 2X Reducing Laemmli Sample Buffer was added to the beads and heated at 100°C for 5 minutes.and supernatant used for Western blot analysis.

### NP-IgG1 ELISA

ELISA plates (675061, Greiner Bio-One) were coated with 5 μg/ml NP-BSA (N-5050H-10, Biosearch Technologies) in bicarbonate/carbonate buffer at 4°C overnight. After three washes with PBS, plates were blocked with 5% skimmed milk in PBS at 37°C for 1 hour, followed by 3 x PBS washes. Serum samples diluted in 1% milk were added and incubated at 37°C for 1 hour. For relative quantification, a standard serum sample was made by pooling samples on day 7, post-secondary immunization from a prior experiment, then aliquoted for all subsequent experiments. Serum samples of various days were serially diluted first to determine the proper dilution, and 1:1000-1:5000 was chosen for NP-IgG1 ELISA. After serum sample incubation, plates were washed 6 times with PBS-0.05% Tween 20 followed with detection antibody incubation at 37°C for 1 hour. Alkaline phosphatase-(AP-) conjugated goat-anti mouse IgG1 (Southern Biotech) detection antibody was diluted in 1% milk-PBS at a ratio of 1:2000. After 5 washes with PBS-0.05% Tween 20 and once with PBS, AP-substrate (S0942, Sigma) dissolved in pNPP buffer was added for 15-20 min and absorbance was measured at 405 nm by ELISA plate reader (FLUOstar Omega, BMG Labtech).

### ELISpot

MultiScreenHTS-HA filter plates (MSHAS4510) were first rinsed with 35% ethanol for 30 seconds and washed 3 times with PBS. Plates were coated with 20 μg/ml NP-BSA in PBS at 4°C over night. After three washes with PBS, plates were blocked with RPMI-1640 medium supplemented with 10% FBS at 37°C for 30 minutes. Then bone marrow cells were added in duplicates in culture medium at the density of 3 × 10^5^ /100 μL/ well and cultured at 37°C over night. Plates were then washed 3 times with PBS and 3 times with PBS-0.05% Tween 20. AP-conjugated anti-mouse IgG1 detection antibody diluted in 1% FBS was added to plates for 1 hour incubation at 37°C. After 5 washes with PBS-0.05% Tween 20 and once with PBS, AP substrate (170-6432, Bio-Rad) was added for spot development. Plates with clear spots and clean background were washed with water to stop development, dried, and counted with the AID ELISpot Reader System (ELR078IFL, AID GmbH).

### Plasmid construction

The TFEB polyproline motif-mCherry-IRES, 13P-mCherry-IRES, Random-mCherry-IRES, murine wildtype/mutant (PPP to AAA) *Tfeb-HA* were cloned into pMSCV-IRES-EGFP with HindIII and BamHI. A flexible GSGSG linker was inserted between the targeted sequence and mCherry to assist folding of the fusion protein. The sequence coding C-terminal 37-amino acid degron of murine ODC (Kelly et al., 2007) was inserted to the C-terminal of mCherry with Not1 and Sal 1.

### Cell drug treatments

GC7 (Millipore) was added for 24 hours to B cells (10 μM) on day 2 after B cell stimulation (unless otherwise indicated) or to Jurkat/NIH 3T3 cells (100 μM). 10 nM bafilomycin A1 (Cayman Chemical), 10 μg/ml cycloheximide (Sigma), or 100 nM Torin 1 (Cayman Chemical) were added to cells for 2 hours. 10 μM etoposide (Cayman Chemical) was added to cell culture for 6 hours, or 1 mM difluoromethylornithine (DFMO, Enzo Life Sciences) for 24 hours.

### Cell culture systems

Cells (Jurkat, NIH 3T3, HEK293T) were cultured in RPMI-1640 medium supplemented with 10% FBS, Penicillin-Streptomycin (P/S), and L-Glutamine (all Sigma) in 5% CO_2_ at 37°C. The IPTG-inducible MISSION shRNA lentiviral vector pLKO-puro-IPTG-3xLacO contains shRNA against the 3’-UTR of murine eIF5A mRNA (custom-made from #SHCLND-NM181582-TRCN0000125229, Sigma) or a corresponding non-target shRNA control (#SHC332-1EA, Sigma). 100 μM IPTG (Thermo Fisher) was used to induce the expression of shRNA. Stable transduction of the lentivirus was performed as previously described (Preukschas et al., 2012). Immortalized *Dohh^flox/flox^* and *Dohh^+/+^* 3T3 cells were established from mouse embryonic fibroblasts (Sievert et al., 2014). 100 nM 4-OHT (H6278, Sigma) was used to induce the knockout of *Dohh*. NIH 3T3 cells were transfected with siRNA according to Thermo Fisher protocol (13778075, Lipofectamine^®^ RNAiMAX Reagent). Cells were collected on day 3 posttransfection for western blot analysis. For plasmid transfection assay, NIH 3T3 cells pretreated with 10 μM GC7 for 24 hours were transfected with the indicated plasmids using Lipofectamine 3000 according to the manufacturer’s protocol. mCherry and GFP expression was quantified by flow cytometry 24 hours post transfection.

### Flow cytometry

Cells were stained with fixable Zombie Aqua Live/Dead staining (423102, Biolegend), FcR block, followed by surface marker antibodies and analyzed with four-laser LSR Fortessa X-20.

Acquired data were analyzed using FlowJo 10.2. For LC3-II flow cytometry (FACS) staining, the FlowCellect^TM^ Autophagy LC3 antibody-based assay kit (FCCH100171, Millipore) and for CellTrace staining, CellTrace Violet (C34557, Thermo Fisher) were used according to the manufacturer’s protocol.

### CytoID staining

Cells were stained with CytoID (ENZ-51031-K200, Enzo Life Sciences) according to manufacturer’s protocol. Briefly, cells were re-suspended in CytoID staining solution (1:4000 diluted in RPMI without phenol red (R1780, Sigma) supplemented with 5% FBS) and incubated at 37°C for 30 minutes in the dark. Then cells were washed once, followed by surface marker staining and FACS analysis without fixation.

### Quantitative PCR

RNA was extracted using the RNeasy Plus Mini Kit (74134, Qiagen). The concentration of RNA was measured using NanoDrop1000 (Thermo Fisher), followed by reverse transcription using the High Capacity RNA-to-cDNA Kit (4387406, Thermo Fisher). Taqman probes (Thermo Fisher), TaqMan Gene Expression Master Mix (4369016, Thermo Fisher), and the ViiA 7 Real-Time PCR System (Thermo Fisher) were used for quantitative PCR. ΔΔCt method was used for the quantification of target mRNAs expression using *Gapdh* as the reference gene.

### Confocal microscopy

B cells from GFP-LC3 mice were MACS-purified and fixed with 4% paraformaldehyde at room temperature for 10 min. After nuclear staining with DAPI (Sigma), cells were transferred to slides using Cytospin 3 cytocentrifuge (Shandon) and imaged with the Olympus FV1200 Laser Scanning Microscope. CellProfiler software was used for automatic autophagosome quantification. Nuclei were defined in DAPI channel (diameter of 45-100 pixel units), autophagosomes were defined in GFP channel (diameter of 3-15 pixel units,1 pixel unit =100 nm). An artificial but consistent threshold of 1-10 cellular GFP spots in GFP-LC3 confocal imaging was used for statistical analysis. 10%-45% B cells are GFP-in GFP-LC3 transgenic mice as assessed by flow cytometry. Therefore cells with 0 GFP spot (either due to no GFP expression or very low autophagy) were excluded from all samples. Cells with more than 10 GFP spots were also excluded from all samples as outliers.

### Cell fractionation

Cells in a 1.5 mL Eppendorf tube were washed once with cold PBS and re-suspended in HLB buffer (10 mM Tris-HCl pH 7.5, 10 mM NaCl, 2.5 mM MgCl2) with protease inhibitors and phosphatase inhibitors, then HLB+2N buffer (HLB+0.4% NP-40, 1N is 0.2% NP-40) was added and carefully mixed. Samples were incubated on ice for 5 minutes and then underlayed with 200 uL HLB+NS buffer (HLB+0.2% NP-40+10% sucrose). Samples were centrifuged at 500 g for 5 minutes and the upper half of the supernatant was collected as the cytoplasmic fraction. The remaining liquid was discarded and the nuclear pellet was washed once more with HLB buffer and lysed with NP-40 lysis buffer. All procedures were performed on ice.

### Protein mass spectrometry

LPS-stimulated mouse B cells were treated with 10 μM GC7 on day 2 for 24 hours. Cells were collected and fractionated for label-free quantitative mass spectrometry (MS) analysis. For SILAC MS, B cells stained with CellTrace Violet (C34557, Thermo Fisher) were cultured in two types of medium: light and heavy. In light medium, SILAC RPMI-1640 medium (89984, Thermo Fisher) supplemented with 1.15 mM L-Arg (Sigma) and 0.22 mM L-Lys (Sigma), 10% dialysed FBS (Sigma), and P/S, L-Glutamine, 50 μM 2-mercaptoethanol, 20 mM HEPES were used. In heavy medium, ^13^C_6_ ^15^N_4_-L-Arg (Arg-10, Silantes) and ^13^C_6_ ^15^N_2_-L-Lys (Lys-8, Silantes) were used instead of the light arginine and lysine respectively. 10 μM GC7 was added to either light or heavy medium in two repeats. After GC7 treatment, dead cells were firstly removed using the Dead Cell Removal Kit (Miltenyi). An identical number of cells that had divided at least 4 times in light or heavy medium were collected by FACS, and mixed for cell fractionation and protein MS analysis.

Peptide samples were prepared using the Filter Aided Sample Preparation (FASP), as previously described (Wisniewski et al., 2009). Briefly, Vivacon 500 filters (Sartorius, VN01H02 10 kDa/VNCT01) were pre-washed with 200 μL 0.1% trifluoroacetic acid in 50% acetone. Lysate samples were loaded to the filter and denatured with 200 μL 8 M urea in 100 mM triethylammonium bicarbonate buffer (TEAB) for 30 minutes at room temperature. Denatured proteins were reduced by 10 mM tris (2-carboxyethyl) phosphine (TCEP) for 30 minutes at room temperature and alkylated with 50 mM chloroacetamide for 30 minutes at room temperature in the dark. Subsequently, 1 μg LysC (Wako) in 150 μL 50 mM TEAB containing 6M urea was added and incubated at 37°C for 4 hours. Then the buffer was diluted to 2 M urea by 50 mM TEAB, followed by adding 0.5 μg trypsin (Promega) overnight at 37°C. Trypsinised samples were centrifuged and the flow-through, containing peptides, was dried and resuspended in 70 μL 10% formic acid (or 5% formic acid and 5% DMSO for SILAC experiment).

Peptides were separated on an Ultimate 3000 UHPLC system (Thermo Fisher) and electrosprayed directly into a QExactive mass spectrometer (Thermo Fisher) The peptides were trapped on a C18 PepMap100 pre-column (300 μm i.d. × 5 mm, 100 Å, Thermo Fisher) using solvent A (0.1% formic acid in water) at a pressure of 500 bar, then separated on an in-house packed analytical column (75 μm i.d. packed with ReproSil-Pur 120 C18-AQ, 1.9 μm, 120 Å, Dr. Maisch GmbH) using a linear gradient (length: 120 minutes, 15% to 35% solvent B (0.1% formic acid in acetonitrile), flow rate: 200 nl/min). Data were acquired in a data-dependent mode (DDA). Full scan MS spectra were acquired in the Orbitrap (scan range 350-1500 m/z, resolution 70000, AGC target 3e6, maximum injection time 50 ms). The 10 most intense peaks were selected for HCD fragmentation at 30% of normalised collision energy at resolution 17500, AGC target 5e4, maximum injection time 120 ms with first fixed mass at 180 *m/z*. Charge exclusion was selected for unassigned and 1+ ions. The dynamic exclusion was set to 20 seconds.

Raw MS data were processed by MaxQuant (version 1.5.0.35i) for peak detection and quantification (Cox and Mann, 2008; Cox et al., 2011). MS spectra were searched against the *Mus musculus* UniProt Reference proteome (retrieved 12/01/17) alongside a list of common contaminants, using the Andromeda search engine with the following search parameters: full tryptic specificity, allowing two missed cleavage sites, fixed modification was set to carbamidomethyl (C) and the variable modification to acetylation (protein N-terminus) and oxidation (M). The search results were filtered to a false discovery rate (FDR) of 0.01 for proteins, peptides and peptide-spectrum matches (PSM). Protein intensity distributions were log2 transformed and median-centred using Perseus (version 1.5.5.3). For the SILAC analysis, missing values were replaced by an estimated background noise value. Proteins without greater-than-background values in both replicates for at least one condition were removed. MA plots were generated using R (version 3.4.2). Reviewed autophagy proteins were searched against the *Mus musculus* UniProt database. Data from the independently analyzed cytoplasmic and nuclear proteins were shown as overlaid MA-plots wherein the log2 average (GC7 + Vehicle) intensity (A) is plotted on x versus the log2 (GC7:Vehicle) fold change (M) plotted on *y*.

### Spermidine measurement by gas chromatography mass spectrometry (GC-MS)

Cells were washed with PBS and the pellet resuspended in lysis buffer (80% methanol + 5% Trifluoroacetic acid) spiked with 2.5 μM 1,7-diaminoheptane (Sigma). The cell suspension, together with acid-washed glass beads (G8772, Sigma), was transferred to a bead beater tube and homogenized in a bead beater (Precellys 24, Bertin Technologies) for four cycles (6500 Hz, 45 s) with 1 minute of ice incubation between each cycle. The homogenized samples were centrifuged at 13,000 g for 20 minutes at 4°C. The supernatant was collected and dried overnight. For chemical derivatization, 200 μL trifluoroacetic anhydride was added to the dried pellet and incubated at 60°C for 1 hour, shaking at 1200 rpm. The derivatization product was dried, resuspended in 30 μL isopropanol and transferred to glass loading-vials. The samples were analyzed using a GCxGC-MS system as described (Yu et al., 2017). The following parameters were used for quantification of the 1D-GC-qMS data: Slope: 1000/min, width: 0.04 s, drift 0/min and T. DBL: 1000 min without any smoothing methods used.

### OPP-Click Assay

Protein synthesis rate was measured using the Click-iT Plus OPP Protein Synthesis Assay (C10456, Thermo Fisher) according to manufacturer’s protocol. Geometric mean fluorescence intensity (normalized to vehicle) was used as an indicator of the relative protein translation rate.

### Ribosome profiling

The sequencing library was prepared according to the reference (Ingolia, 2010; Ingolia et al., 2013) with the following changes. LPS-activated B cells were MACS-purified with the Dead Cell Removal Kit (Miltenyi) to remove dead cells prior to cell lysis. 100 μg/ml cycloheximide was used in the lysis buffer. The B cell lysate was spiked with the *S. cerevisiae* lysate at the ratio of 13 million B cells: 270 million yeast cells (which equals to 20:4 μg miRNA). Clarified B cell and yeast lysates were RNase I-digested, ribosome purified and miRNA extracted to determine the miRNA yield for spiking. Size exclusion chromatography (27-5140-01, GE Healthcare) was used to recover ribosomes. The miRNeasy Micro Kit (Qiagen) was used to purify miRNAs. The Ribo-Zero Gold rRNA Removal Kits (illumina, MRZG126, MRZY1306) were used to deplete ribosomal RNAs, in which the Ribo-Zero Removal Solutions from the mouse and yeast kits were mixed at the ratio of 8 μL : 4 μL to remove both mouse and yeast rRNAs. The following barcoding primers were used for next generation sequencing. 1. 5’-AATGATACGGCGACCACCGAGATCTACAC GATCGGAAGAGCACACGTCTGAACTCC AGTCAC<u>ATGCCA</u>TCCGACGATCATTGATGG. 2. 5’-AATGATACGGCGACCACCGAGATCTACAC GATCGGAAGAGCACACGTCTGAACTCC AGTCAC<u>TGCATC</u>TCCGACGATCATTGATGG. The libraries were sequenced at the Biopolymers Facility, Harvard Medical School.

For the purposes of alignment, the reference genome assemblies for Mus musculus (GRCM38) and Saccharomyces cerevisiae (SacCer3) were concatenated to create a combined genome fasta file. A combined rRNA transcript file was also created using the sequences listed in Table S1. Prior to alignment, Adaptor sequence (ATCTCGTATGCCGTCTTCTGCTTG) was removed from demultiplexed reads and low quality reads (Phred score < 20) were discarded. To remove contaminating rDNA reads, reads were aligned to the combined rDNA genome using bowtie2 and all unaligned reads were kept for alignment to the combined genome. Non-rDNA reads were aligned to the combined genome using bowtie2 and the following parameters: -t -N 1 and uniquely aligning reads were extracted using samtools view -bq 10.

All post-alignment analysis was performed using custom scripts written in R. P-site offsets for each read-length from 26 to 35 nts in the rpf libraries were determined by generating metagenes relative to the START codon and applied to aligned reads (Table S2). For all metagenes, transcripts with no reads within the window of interest were excluded from analysis before applying any other filters. Additionally, for each position in the windows of the 5’ and 3’ metagenes, the highest and lowest 0.1% of signals were removed from analysis.

The 100 most stalled PPP motifs were identified using the “stall-ratio” defined as the ratio of coverage over position +3 relative to the first proline against the mean coverage of the entire 80 bp window. The following filters were also applied to exclude windows with only a few sporadic spikes which may not reflect real translation. To be considered, a window must contain > 10 reads outside of the +3 position and the window must have reads in at least 5 positions. After applying these filters, the 100 genes with the highest stall-ratios were selected for metagenes.

### Human B cell assays

25 ml of blood was obtained from non-fasting healthy donors, old (≥ 65 y of age) young (≤65y of age) in heparinized tubes. All subjects gave written informed consent. The study was approved by the Ethics Committee of Oxford University. Peripheral blood mononuclear cells (PBMCs) were isolated by density gradient centrifugation using Ficoll-Paque. PBMCs were counted using trypan blue. They were either freshly used or were frozen until further use. Thawed cells were placed in full medium overnight to allow selection of viable cells. PBMCs were washed with PBS and lysed using NP-40 lysis buffer containing proteinase inhibitors (Sigma) and phosphatase inhibitors (Sigma) on ice for Western blot assay or using in lysis buffer (80% methanol + 5% Trifluoroacetic acid) spiked with 2.5 μM 1,7-diaminoheptane (Sigma) for Spermidine measurement by GC-MS. B cells were sorted using a negative selection kit (B Cell Isolation Kit II, human, Miltenyi Biotec) according to the manufacturer’s protocol. B cells (5 × 105 cells) were seeded in 24-well plates, then activated with anti-IgM (5 μg/ml, Jackson Immuno Research) and CD40L (100 ng/ml, Enzo Life science) and treated with 1 mM DFMO, 10 μM spermidine, or with both for 7 days. Cells then were lysed for western blotting or stained for LC3 measurement by flow cytometry. IgG release in culture supernatants was measured by heterologous two-site sandwich ELISA, according to the manufacturer’s protocol (Invitrogen).

### Statistical analyses

Prism software (GraphPad) was used for statistical analyses. Data are represented as either box-and-whisker plots showing whiskers and outliers by Tukey method or mean ± SEM. All data points refer to the measurement of distinct samples (biological repeats). Paired or unpaired twotailed Student’s t-test was used for comparisons between two normally distributed data sets with equal variances. One-tailed Student’s t-test was used when the null hypothesis has a clear direction according to current knowledge (*e.g*. spermidine is an autophagy inducer in multiple systems or Bafilomycin A1 treatment leads to accumulation of LC3-II). Welch’s unequal variances t-test was used when the two normally distributed data sets have unequal variances. Holm-Sidak method was used to adjust P values of a family of multiple t-tests with a single null hypothesis. Paired or unpaired one/two-way ANOVA, followed with post hoc Sidak’s test (comparing two means within each group), Dunnett’s test (comparing multiple means with the single control within each group), or Tukey’s test (comparing the mean of each sample with the mean of every other sample), was used for multiple comparisons of normally distributed datasets with one/two variables. Mann-Whitney U test was used for comparisons between non-parametric datasets. Linear regression with 95% confidence interval was used to assess the relationships between age and the expression of target proteins or spermidine levels, in which R^2^ was used to assess the goodness of fit and the P value calculated from F test was used to assess if the slope was significantly non-zero. P value was used to quantify the statistical significance of the null hypothesis testing. *P≤0.05, **P≤0.01, ***P≤0.001, ****P≤0.0001.

## ACKNOWLEDGMENTS

We thank the staff of the University of Oxford Biomedical Services Unit for animal care, Jonathan Webber and Craig Waugh for assistance with FACS sorting, and Volodymyr Nechyporuk-Zloy for assistance with confocal microscopy imaging, Alfredo Castello Palomares for help with SILAC proteomics and providing general advice. Alexander Clarke is acknowledged for reading the manuscript. H.Z. is funded by the China Scholarship Council and Elysium Health fellowship and the A.K.S. lab is funded by a Wellcome Trust Investigator Award (103830/Z/14/Z). S.B. is funded by a Swiss National Science Foundation Grant (31003A_150066). J.F. is funded by the Wellcome Trust Chromosome and Developmental Biology PhD Program (ALR00520). J.M. lab is funded by BBSRC grant (BB/P00296X/1). Li-Cor Odyssey imager is funded by ERC (AdG 670930).

## AUTHOR CONTRIBUTIONS

HZ designed and performed most experiments, GA performed human B cell studies, JF helped to set up and analyze ribosome profiling in primary B cells, PC and SM generated proteomic data, YS/LF ran some Western blots, SB provided KD/KO cell lines and Dhs KO bone marrow, TR helped to set up the Dhs KO chimera experiments, JM and SB gave advice on data interpretation and read manuscript, AKS supervised this project (HZ’s thesis), designed the experiments, provided funds, and wrote the manuscript.

There are no financial competing interests.

---

**Figure S1.**
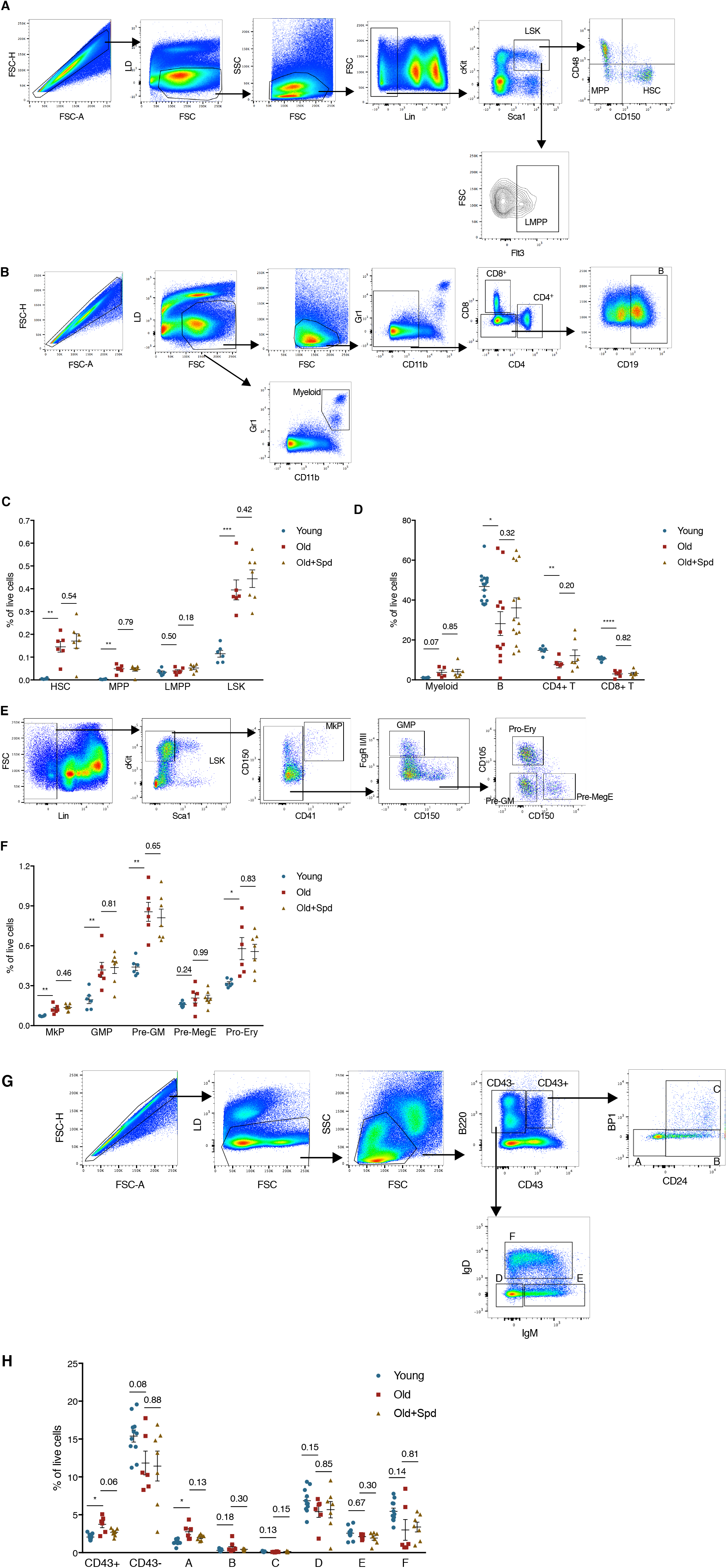
Related to Figure 1. Spermidine does not Affect Hematopoiesis in Old Mice. Various hematopoietic cell types in bone marrow and spleen from young (12 weeks), old (22-24 months), and old mice continuously administered with spermidine in drinking water for 6 weeks as in Fig. 1A were assessed by flow cytometry. (A) Gating strategy for hematopoietic stem cells (HSCs), multipotent progenitors (MPPs), lymphoid-biased multipotent progenitors (LMPPs), and Lin^-^Sca1^+^cKit^+^ cells (LSKs, enriched of hematopoietic stem and progenitor cells) in bone marrow. (B) Gating strategy for myeloid cells, B cells, CD4^+^ T cells, and CD8^+^ T cells in spleen. (C) Expanded phenotypic HSCs, MPPs, and LSKs in bone marrow from old mice. The abundance of indicated cell types as % of total live cells is shown. n = 6-7 mice as indicated by dots. (D) Spleen lineages are lymphopenic in old mice. n = 6-17 mice as indicated by dots, combined from 2 independent experiments. (E/F) Expanded myeloid progenitors in bone marrow from old mice. (E) Gating strategy for megakaryocyte progenitors (MkPs), granulocyte-macrophage progenitors (GMPs), pre-granulocyte/macrophages (Pre-GMs), pre-megakaryocytes/erythrocytes (pre-MegEs), and pro-erythroblast cells (Pro-Erys). (F) The abundance of indicated cell types as % of total live cells. n = 6-7 mice as indicated by dots. (G/H) Accumulated pro-B cells in old mice. (G) Gating strategy for Hardy fractions A-F. (H) The abundance of indicated cell types as % of total live cells. n = 7-11 mice as indicated by dots, pooled from 2 independent experiments. Data represented as mean ± SEM. Two-tailed Welch’s t-test. *P≤0.05, **P≤0.01, ***P≤0.001, ****P≤0.0001.

**Figure S2.**
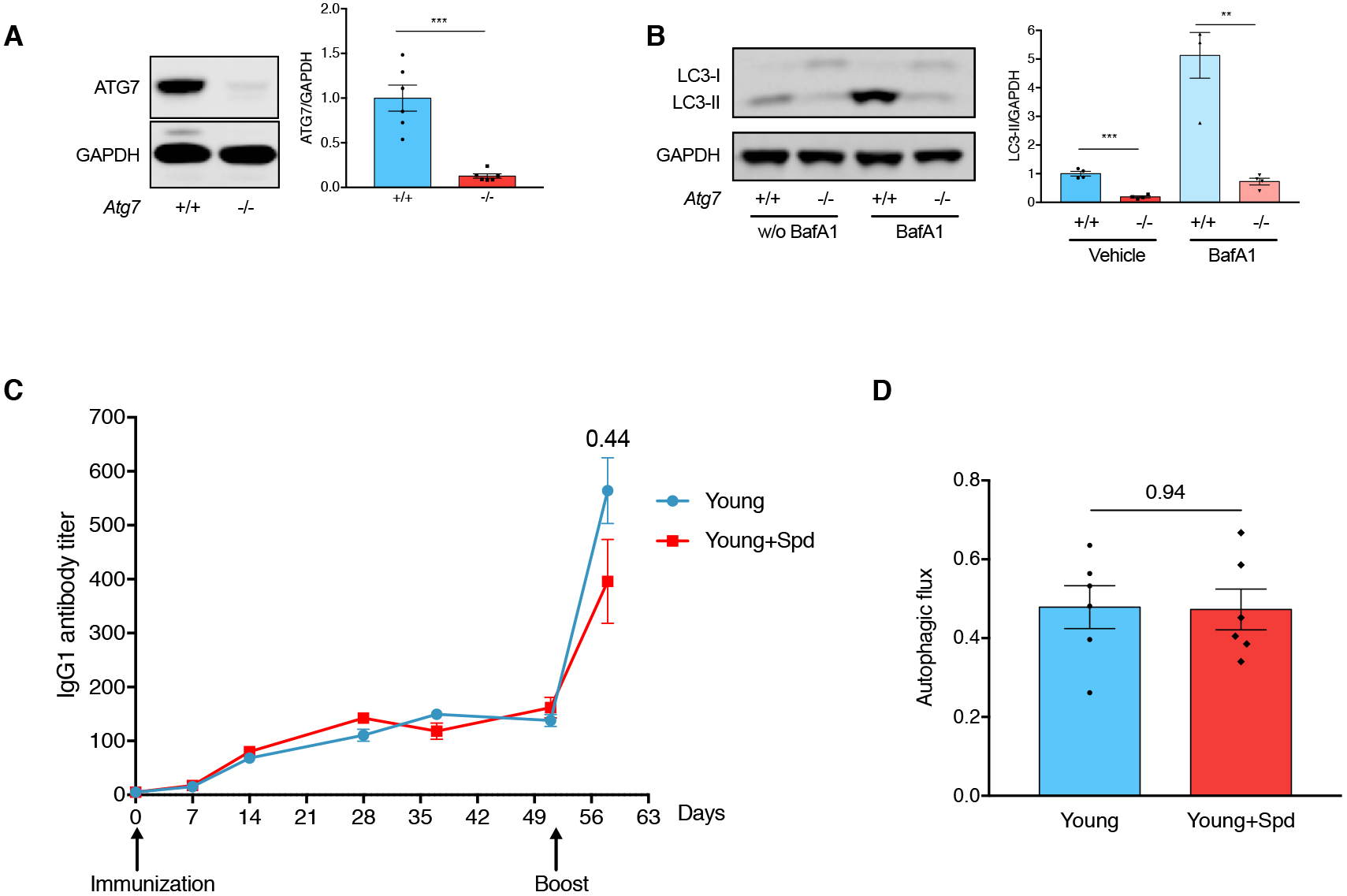
Related to Figure 2. Spermidine does not Improve B Cell Responses in Young Mice. (A/B) Confirmation of reduced ATG7 expression and autophagy in B cells purified from B cell-specific *Atg7*-knockout mice. (A) B cells were purified from *Mb1-Cre-, Atg7^flox/flox^*(+/+) and *Mb1-Cre+, Atg/^flox/flox^* (-/-) mice and the expression of ATG7 was assessed by Western blot (left). For quantification (right), n = 6 mice. (B) Purified B cells in (A) were cultured with 10 nM BafA1 for 2 h for LC3-II measurement by Western blot (left). For quantification (right), the LC3-II bands intensity was normalized to GAPDH first and then to the average of Vehicle +/+. n = 4 mice. (C/D) Spermidine does not improve IgG1 responses in young mice. (C) Young adult mice were immunized and boosted with NP-CGG. Spermidine administration and serum NP-specific IgG1 measurement were processed as in Fig. 2D. n = 11 (D7), 19 (D14), 12 (D28), 9 (D37), 15 (D51) and 6 (D58) mice, combined from 3 independent experiments. (D) Autophagic flux of splenic B cells (B220+CD19+) from mice culled on D58 in (C) was measured by LC3-II flow cytometry staining as in Fig. 1 A. n = 6. Data represented as mean ± SEM. One-tailed Welch’s t-test (A/B). Two-tailed Student’s t-test (C/D). **P≤0.01, ***P≤0.001.

**Figure S3.**
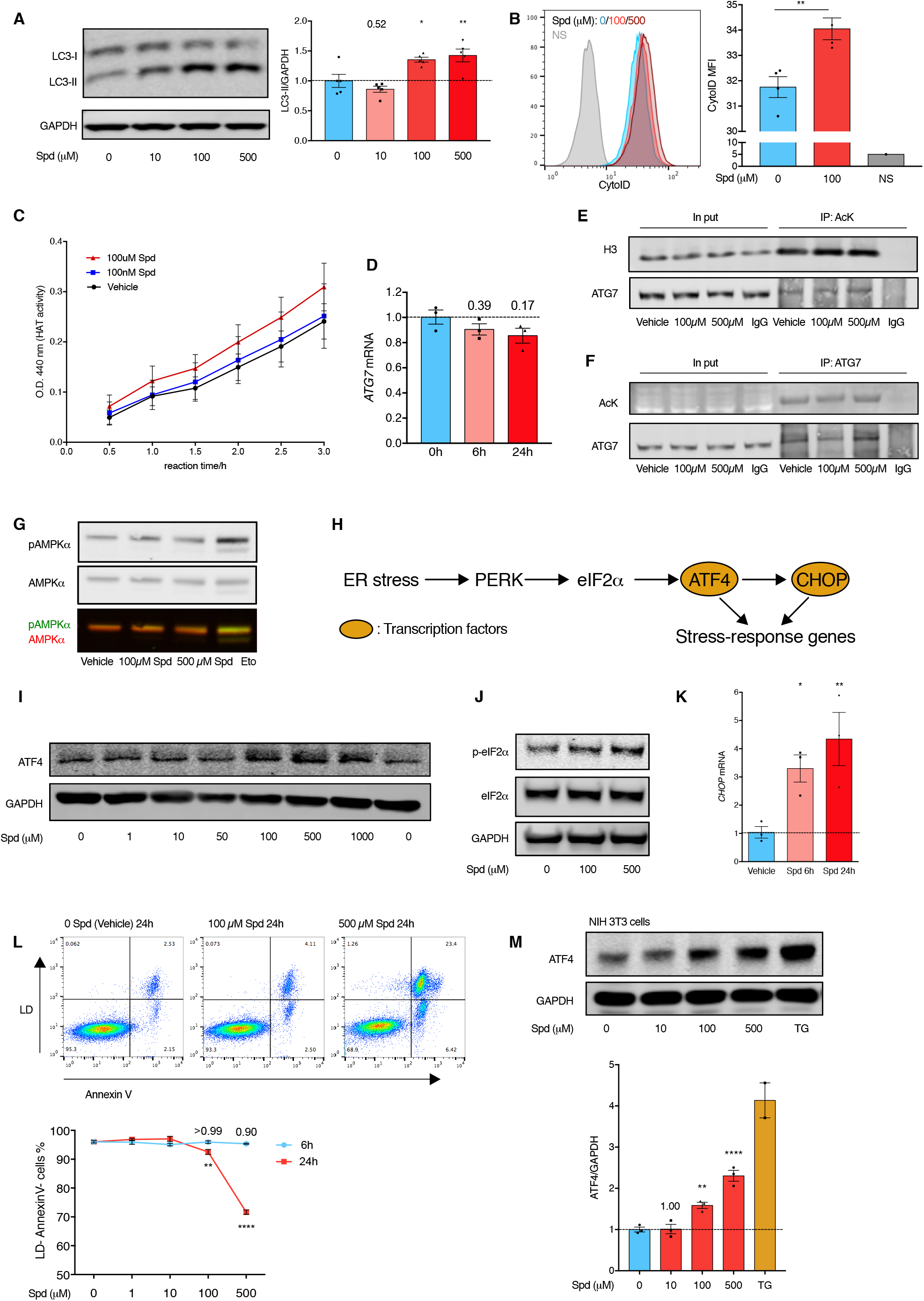
Related to Figure 3. Spermidine does not Directly Inhibit HAT Activities and High-dose Spermidine Induces Cellular Stress. (A/B) Spermidine induces autophagy *in vitro*. Jurkat cells were treated with spermidine (Spd) as indicated for 6 h. Autophagy was assessed by LC3-II Western blot (A). n = 5. Autophagosome/autolysosome-specific staining of CytoID was measured by flow cytometry (B). n = 4. (C-F) Spermidine does not inhibit HAT activity in Jurkat cells. (C) Jurkat nuclear extract was prepared using the Nuclear/Cytosol Fractionation Kit, incubated with spermidine of indicated concentrations (starting from 0h) and the relative HAT activity was measured using the HAT Activity Colorimetric Assay Kit. n = 3. (D) Jurkat cells were treated with 100 *μ*M spermidine for 6h or 24h. *ATG7* mRNA was measured by quantitative PCR (qPCR) with *GAPDH* used as the reference gene. n = 3. (E/F) Jurkat cells were treated with spermidine for 6h and cellular proteins with acetylated lysine residues were pulled down (IP: AcK) and assessed for H3 and ATG7 acetylation (E). To assess ATG7 acetylation in an alternative way, ATG7 was pulled down (IP: ATG7) and acetylation measured with an antibody against AcK (F). (G) Spermidine does not affect AMPK activity. Jurkat cells were treated with spermidine of indicated concentrations for 6 h. AMPK activity was assessed by AMPKα phosphorylation. Cells were treated with 10 *μ*M etoposide for 6h as the positive control. Representative of 3 independent repeats. (H-M) High-dose spermidine induces cellular stress. (H) Schematic overview of the ER stress PERK-eIF2α-ATF4-CHOP pathway. ER stress activates the protein kinase R-like endoplasmic reticulum kinase (PERK), which phosphorylates eIF2α. Phosphorylated eIF2α facilitates the translation of the transcription factor ATF4, which then induces the expression of CHOP. ATF4 and CHOP induce the expression of multiple stress-response genes including chaperones, apoptosis, and autophagy. (I/J) Jurkat cells were treated with spermidine of indicated concentrations for 6h. The expression of ER stress markers ATF4 (I) and phosphorylation of eIF2α (J) were assessed by Western blot. Representative of 3 independent repeats. (K) Jurkat cells were treated with 100 *μ*M spermidine for 6 h or 24 h. The expression of *CHOP* was assessed by qPCR with *GAPDH* as the reference gene. n = 3. (L) Jurkat cells were treated with spermidine of indicated concentrations for 6 h or 24 h. Cell viability and apoptosis were assessed by Live-Dead (LD) and Annexin V flow cytometry staining. n = 3. (M) NIH 3T3 cells were treated with spermidine of indicated concentrations or thapsigargin (TG) for 6 h. The expression of ATF4 was assessed by Western blot. n = 3. One-way ANOVA with post hoc Dunnett’s test comparing with 0 *μ*M/Vehicle (A/K/M) or 0 h (D). Student’s t-test (B). Two-way ANOVA with post hoc Dunnett’s test comparing with 0 *μ*M (L). Data represented as mean ± SEM. *P≤0.05, **P≤0.01, ****P≤0.0001.

**Figure S4.**
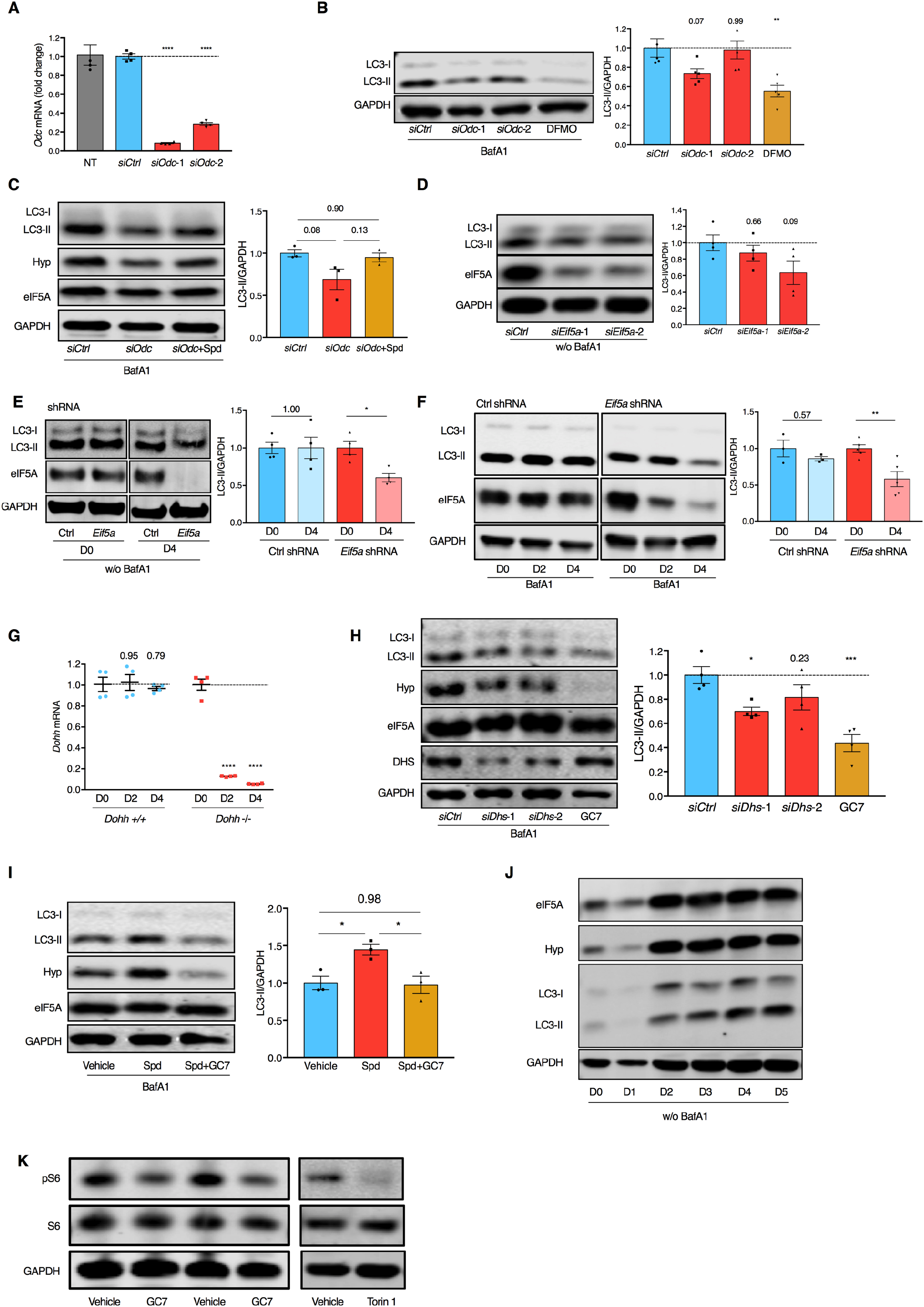
Related to Figure 3. Spermidine Maintains Cellular Autophagy by Hypusinating eIF5A. *Odc* knockdown efficiency test. NIH 3T3 cells were transfected with non-targeting siRNA *(siCtrl)* or siRNA targeting two different regions of *Odc* mRNA *(siOdc-1/2)* for 3 days. The expression of *Odc* was assessed by qPCR with *Gapdh* as the reference gene (normalized to *siCtrl)*. n = 4. NT, non-treatment. *siOdc-1* was used in all other figures unless specified otherwise. (B) NIH 3T3 cells were transfected with *siCtrl, siOdc-1*, or *siOdc-2* for 3 days or treated with DFMO for 24 h. LC3-II was measured by Western blot (left) and quantified by normalizing to GAPDH first and then to the average of *siCtrl* (right, n = 5). (C) NIH 3T3 cells were transfected with *siCtrl* or *siOdc* and treated with 10 *μ*M spermidine for 3 days where indicated. LC3-II expression was measured by Western blot. n = 3. (D) NIH 3T3 cells were transfected with *siCtrl* or siEif5a-1/2 for 3 days. LC3-II was measured by Western blot. n = 4. (E/F) The knockdown of *Eif5a* was induced by 100 *μ*M IPTG in NIH 3T3 cells expressing IPTG-inducible *Eif5a* shRNA. The expression of LC3-II on indicated days post IPTG induction was measured by Western blot (E, without BafA1; F, with BafA1). n = 3-5 as indicated by dots. (G) The expression of *Dohh* on indicated days post 4-OHT induced deletion was assessed by qPCR with *Gapdh* as the reference gene. n = 4. (H) NIH 3T3 cells were transfected with *siCtrl* or siRNA targeting two different regions of *Dhs* mRNA (siDhs-1/2) for 3 days or treated with 10 *μ*M GC7 for 24 h. LC3-II was measured by Western blot. n = 4. *siDhs-1* was used in all other figures unless specified otherwise. (I) Spermidine-depleted NIH 3T3 cells by DFMO treatment were rescued with 10 μM spermidine alone or spermidine together with GC for 24 h. LC3-II was measured by Western blot. n = 3. (J) Purified murine B cells were cultured with LPS for indicated days as in Fig. 3H. The expression of overall eIF5A, hypusinated eIF5A, and LC3-II (without BafA1 treatment) was assessed by Western blot. Representative of 4 independent repeats. (K) GC7 inhibits autophagy in an mTOR-independent manner (not via activating mTOR). LPS-activated murine B cells were treated with 10 *μ*M GC7 for 24 h as in Fig. 3I, or with the mTOR inhibitor Torin 1 for 2 h. The expression of S6 and its phosphorylation (Ser235/236) downstream of mTOR were assessed by Western blot. Representative of 5 independent repeats. BafA1 was added 2 h before harvest where indicated for autophagy measurement. Data represented as mean ± SEM. One-way ANOVA with post hoc Dunnett’s test comparing with *siCtrl* (A/B/D/H), or with post hoc Tukey’s test (C/I). Two-way ANOVA with post hoc Sidak’s test (E/F) or Dunnett’s test (G). *P≤0.05, **P≤0.01, ***P≤0.001, ****P≤0.0001.

**Figure S5.**
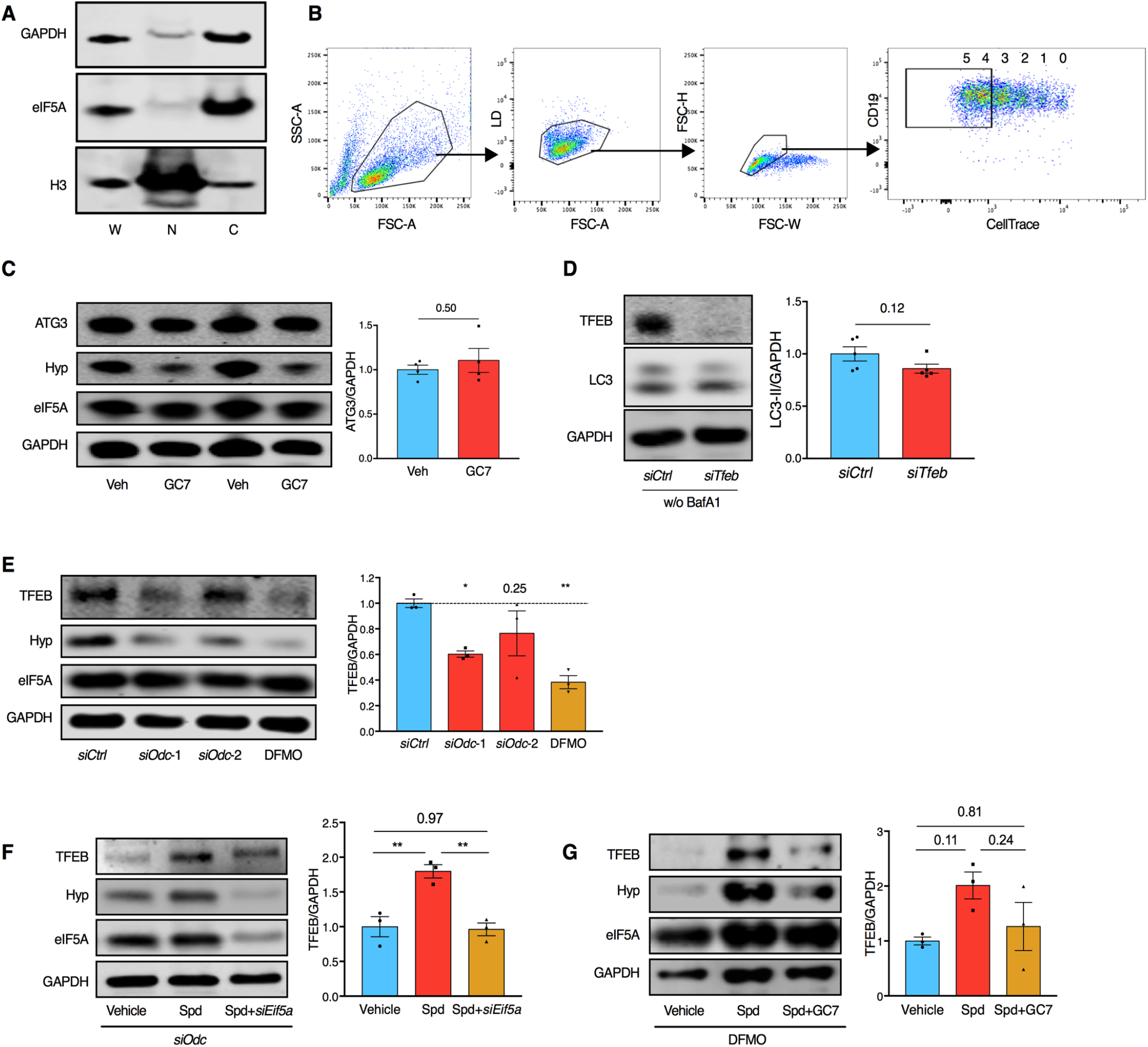
Related to Figure 4. eIF5A Hypusination is Required for TFEB Expression. A. An aliquot of protein samples for MS in Fig. 4A was assessed by Western blot for cell fractionation efficiency. W: whole cell lysate; N: nuclear fraction; C: cytoplasmic fraction.
B. Gating strategy for the FACS sorting in Fig. 4B. Live B cells (CD19^+^) that divided four times or more were collected.
C. LPS-stimulated murine B cells were treated with 10 *μ*M GC7 for 24 h as in Fig. 3I. The expression of ATG3 was assessed by Western blot. n = 4.
D. NIH 3T3 cells were transfected with *siTfeb* for 3 days. LC3-II expression was measured by Western blot. n = 5.
E. NIH 3T3 cells were transfected with *siOdc-1/2* for 3 days or treated with 1 mM DFMO for 24 h. TFEB expression was measured by Western blot. n = 3.
F. *s/Odc*-transfected NIH 3T3 cells were treated (rescued) with 10 *μ*M spermidine and transfected with *siEif5a*. TFEB expression was analyzed by Western blot. n = 3.
G. DFMO-treated NIH 3T3 cells were treated (rescued) with spermidine alone or spermidine together with GC7 for 24 h. TFEB was measured by Western blot. n = 3. Data represented as mean ± SEM. Student’s t-test (C/D). One-way ANOVA with post hoc Dunnett’s test comparing with *siCtrl* (E) or Tukey’s test (F/G). *P≤0.05, **P≤0.01.

**Figure S6.**
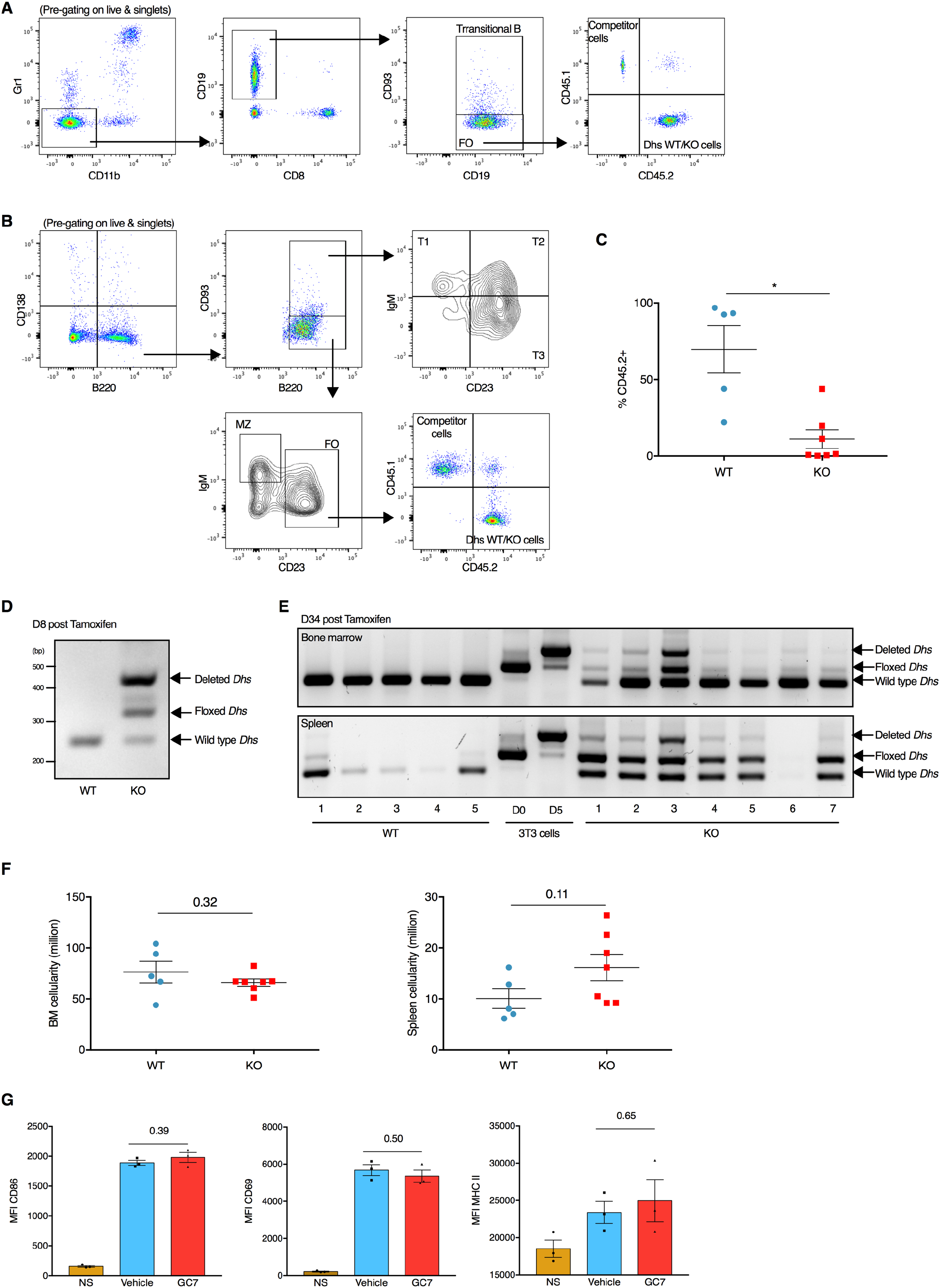
Related to Figure 6. Hypusination of elF5A is Essential for B Cell Development and Activation. (A/B) Gating strategy for transitional (T) B cells, follicular (FO) B cells, marginal zone (MZ) B cells and the contribution of CD45.2^+^ fraction in peripheral blood (A) and spleen (B). (C) The contribution of CD45.2^+^ cells to spleen myeloid cells (CD11b^+^Gr1^+^) of *Dhs* chimera mice in Fig. 6A. n = 5-7 mice as indicated by dots. (D/E) *Dhs* deletion efficiency was tested by PCR. The wild type *Dhs*, floxed *Dhs*, and tamoxifen-induced deletion of *Dhs* were assessed by genotyping DNA extracted from pooled peripheral blood of the chimeric mice on day 8 post tamoxifen administration (D) or from bone marrow and spleen on day 34 post tamoxifen administration (E). 4-OHT-induced *Dhs* deletion (D0/D5) in 3T3 cells derived from CAG-Cre/Esr1^+^, *Dhs^f/f^* mouse embryonic fibroblasts was used as positive control. (F) The cellularity of bone marrow (tibia and femur of both sides) (left) and spleen (right) of WT and KO *Dhs* chimera mice in Fig. 6A. n = 5-7 mice as indicated by dots. Student’s t-test. (G) Quantification of Fig. 6F. n = 3 mice. Data represented as mean ± SEM. Welch’s t-test (C). Student’s t-test (F/G). *P≤0.05.

**Figure S7.**
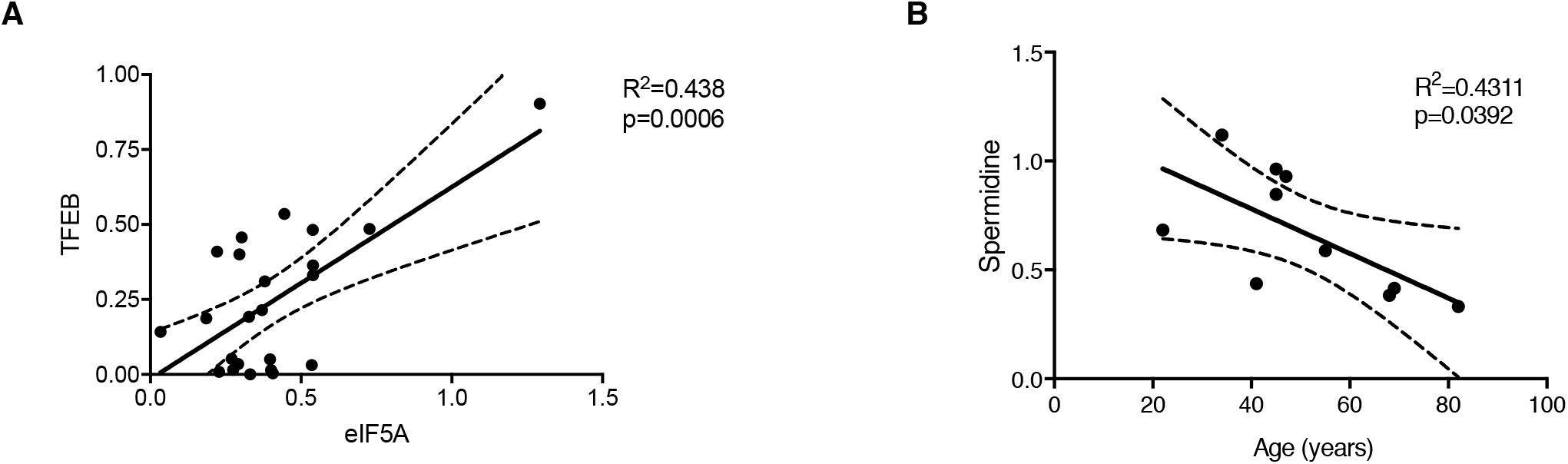
Related to Figure 7. Spermidine Declines with Age in Human PBMCs. A. Correlative analysis between the expression of TFEB and eIF5A (both normalized to GAPDH) in samples of Fig. 7A. n = 23 donors.
B. Spermidine content of PBMCs collected from healthy donors was measured by GC-MS. n = 10 donors. Linear regression with 95% confidence intervals. The goodness of fit was assessed by R^2^. The P value of the slope (whether significantly non-zero) is calculated by F test.

